# Transposable element accumulation drives size differences among polymorphic Y chromosomes in Drosophila

**DOI:** 10.1101/2021.06.02.446622

**Authors:** Alison H. Nguyen, Doris Bachtrog

## Abstract

Y chromosomes of many species are gene poor and show low levels of nucleotide variation, yet often display high amounts of structural diversity. Dobzhansky cataloged several morphologically distinct Y chromosomes in *Drosophila pseudoobscura* that differ in size and shape, but the molecular causes of their dramatic size differences are unclear. Here we use cytogenetics and long-read sequencing to study the sequence content of polymorphic Y chromosomes in *D. pseudoobscura*. We show that Y chromosomes differ by almost 2-fold in size, ranging from 30 to 60 Mb. Most of this size difference is caused by a handful of active transposable elements (TEs) that have recently expanded on the largest Y chromosome, with different elements being responsible for Y expansion on differently sized *D. pseudoobscura* Y’s. We show that Y chromosomes differ in their heterochromatin enrichment, expression of Y-enriched TEs, and also influence expression of dozens of autosomal and X-linked genes. Intriguingly, the same helitron element that showed the most drastic amplification on the largest Y in *D. pseudoobscura* independently amplified on a polymorphic large Y chromosome in *D. affinis*, suggesting that some TEs are inherently more prone to become deregulated on Y chromosomes.

## Introduction

Y chromosomes are a fascinating part of the genome in species with heteromorphic sex chromosomes (1). Not only do Y chromosomes determine maleness in many taxa (2, 3), but their clonal and male-limited transmission is responsible for several unique evolutionary processes that shape their genomic composition (4, 5). Y chromosomes are derived from ordinary autosomes, but over time they often have diverged considerably from their former homolog, the X chromosome (2, 6).

Y chromosomes are transmitted through males only, which makes them an ideal location for male-beneficial genes (7). However, Y chromosomes also have a reduced effective population size and lack recombination; this dramatically decreases the efficacy of natural selection and results in the erosion of ancestral genes on the Y (2, 6). Y chromosomes often accumulate repetitive DNA and evolve a heterochromatic appearance (8). Indeed, old Y chromosomes of many species are typically regarded as repeat-rich genetic wastelands.

A lack of recombination and the small effective population size implies that variation should be low on Y chromosomes, and levels of nucleotide diversity are indeed highly reduced on Y chromosomes of various species (9, 10). Yet, polymorphic Y chromosomes in Drosophila influence a variety of traits, including male fertility (11), temperature tolerance (12), lifespan (13) and expression of 100s of genes across the genome (14, 15). It has been speculated that structural variation involving repetitive regions influences these traits, by globally modulating the genome-wide balance of heterochromatin within the genome (16, 17). Yet, the molecular basis of structural Y variation has not been established.

Morphologically distinct Y chromosomes that occur in natural *D. pseudoobscura* males have first been described almost a century ago (18), and Dobzhansky categorized seven morphologically distinct Y chromosomes that differ in size and shape (19). The molecular basis of size variations of *D. pseudoobscura* Y chromosomes and their origins is unknown, but Dobzhansky speculated that polymorphic Y chromosomes derived by losses of sections from the largest Y (19). He also concluded that morphological and physiological characteristics of a male are not affected by the type of its Y chromosome.

Here, we take advantage of recent advances in sequencing technology, to revisit the dramatic difference in Y size among *D. pseudoobscura* strains. We combine cytogenetic techniques with whole-genome sequencing and chromatin and transcriptome profiling to study the sequence composition of different Y chromosomes, and their phenotypic consequences. We show that mobilization of a handful of TEs and expansion of a satellite sequence (the intergenic spacer region of the rDNA repeats) contributed to size differences among polymorphic Y chromosomes, and we detect subtle differences in heterochromatin enrichment, expression of Y-enriched TEs, and expression of dozens of autosomal and X-linked genes among males containing different Y chromosomes.

## Results & Discussion

### Cytogenetic approaches to identify and characterize different Y chromosomes

We generated Y replacement lines, in order to avoid confounding genomic background effects when contrasting different Y chromosomes (20). We crossed males from different localities with virgins from the sequenced reference strain of *D. pseudoobscura* (MV25), and repeatedly backcrossed males resulting from this cross with virgin females of the reference strain for 9 generations (**Figure S1, S2**). This crossing scheme should ensure that resulting fly lines contain different Y chromosomes in an otherwise isogenic background, that is, all differences in sequence composition among lines should be attributed to the Y chromosome. A total of 26 Y-replacement lines were generated, from diverse geographic populations (**Table S1**, **Figure S1-S3**). To identify Y chromosomes that differ in their morphology and their overall sequence content, we resorted to classical cytogenetic approaches. We first used microscopy to broadly classify Y’s that differ in shape and size from natural strains of *D. pseudoobscura*, (**Figure 1A, Table S2, Figure S3**). In particular, we measured total Y size (relative to the X), and Y shape (long vs. short arm length) to identify different Y types (**Table S2, Figure 1B**). Consistent with Dobzhansky’s studies (19), we find dramatic variation in Y size and shape: we found 22 acrocentric, 3 submetacentric, and 1 metacentric Y chromosome (**Figure 1B, Table S3**).

**Figure 1.**
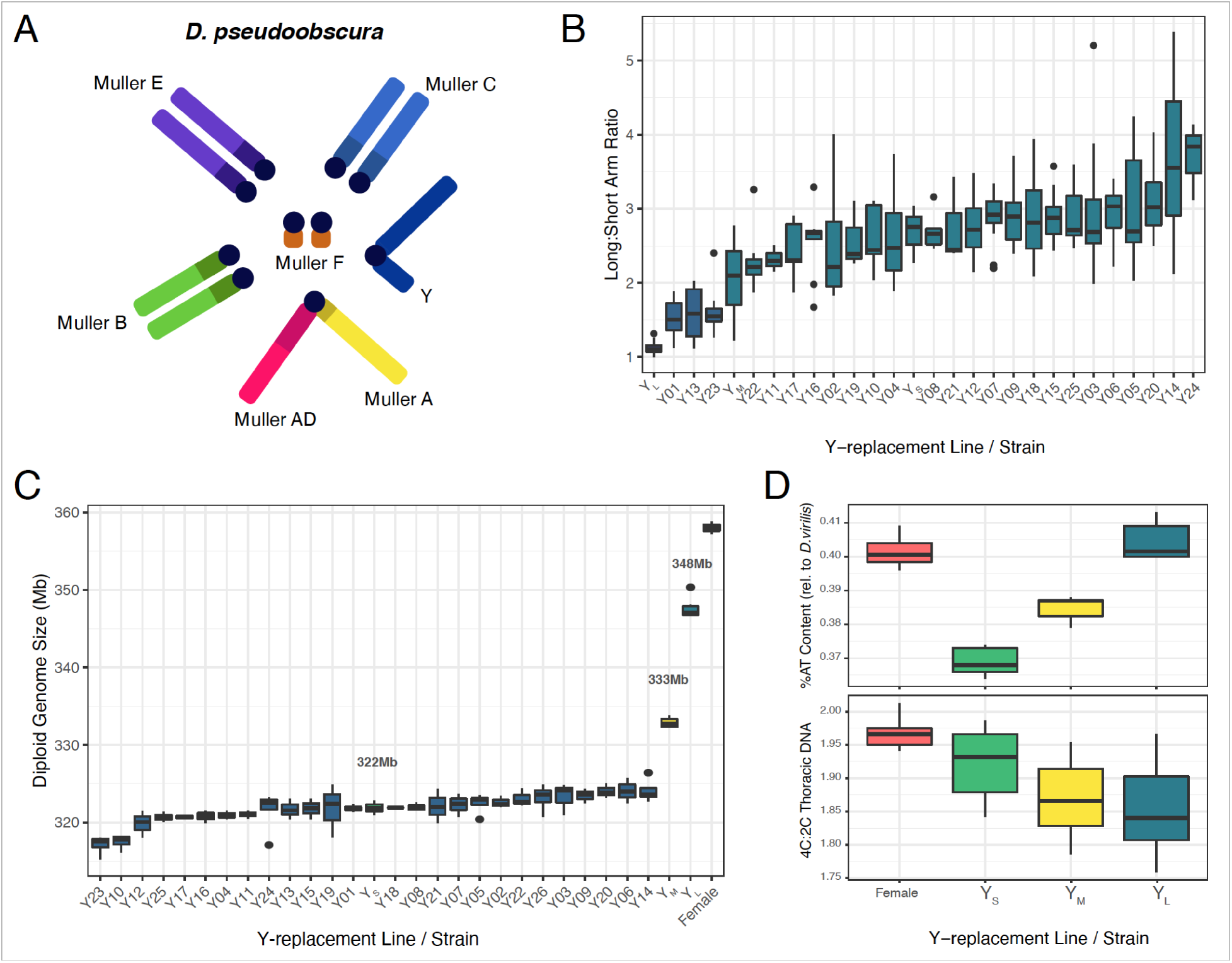
*D. pseudoobscura* male karyotype and Y chromosome size variation. **A.** *D. pseudoobscura* male karyotype. **B.** Measurements of Y chromosome arms (long, short) from chromosome spreads for each Y-replacement line. **C.** Diploid genome size estimations of Y-replacement line males from flow cytometry. Rightmost boxplot (red) is the diploid genome size of the sequenced/backcross female. **D.** Estimates of heterochromatin in 3 Y-replacement line males by staining whole nuclei with DAPI (top) and by staining thoracic cells with propidium iodide (bottom).

To further characterize the size of these Y’s, we used flow cytometry to estimate the relative size of the Y against a *D. virilis* standard (**Figure 1C, Table S4**). We found that inferred diploid genome sizes vary drastically among different Y replacement lines, ranging from 318 to 348 Mb. Assuming that the rest of the genome is roughly 291 Mb (21), this implies that the absolute size of the Y chromosomes vary from 27 to 57 Mb.

Most of this size difference is presumably due to differences in repetitive heterochromatin among Y chromosomes. We used two cytometric approaches to test if males with larger Y chromosomes harbor more heterochromatin. First, we estimated nuclear DNA of cells known to under-replicate heterochromatin. *Drosophila* thoracic cells exhibit endoreplication, the process in which DNA is synthesized in S phase but mitosis does not immediately follow, and the DNA in these cells under-replicate heterochromatin (22). We therefore compared the amount of DNA in G1-phase cells to endoreplicated cells with propidium iodide to obtain proxy estimates for heterochromatin content across Y-replacement males. As a control, we compared the amount of DNA in G1-phase and G2-phase cells from brain nuclei, which replicate all DNA. We found that the largest Y-replacement strain male had a higher proportion of DNA under-replicated in thoracic cells compared to the sequenced strain male and female (**Figure 1D**, **Table S5**).

Heterochromatin is generally AT-rich in contrast to GC-rich euchromatin (23). Therefore, we also used DAPI, a stain that preferentially binds to AT sequences, to obtain independent estimates on heterochromatin content (24). We found that both proxy measurements for heterochromatin agreed well with each other; that is, more under-replicated DNA in thoracic cells corresponds to higher %AT content (**Figure 1D**, **Table S5**). Thus, our cytogenetic work suggests that naturally isolated Y chromosomes in *D. pseudoobscura* differ up to almost 2-fold in size, presumably driven mostly by different repeat composition (i.e. heterochromatin).

### Assembly of a small, medium and large Y chromosome

We chose three Y chromosomes that spanned the entire spectrum of sizes for further molecular characterization. In particular, we selected a small, medium and large Y, and we refer to them as Y_S_, Y_M_ and Y_L_. We obtained high-coverage Illumina sequencing reads (15-19x coverage, obtained by standard PCR+ libraries done on gDNA from head tissue) for each strain, as well as long-read Nanopore sequencing (**Table S6**). Nanopore reads were used for *de novo* assembly of the different Y chromosomes. Assemblies of repetitive regions, including the Y, are challenging and these regions are often collapsed, highly fragmented, or missing in whole-genome assemblies (25, 26). We thus followed a previously described procedure to obtain heterochromatin-enriched assemblies, which were shown to perform superior to recover Y-linked fragments in *D. melanogaster* (26). We used two different assembly algorithms (27, 28), and merged the resulting assemblies (29), in order to produce a more contiguous and complete assembly (**Table S7**). Y-linkage of contigs was inferred using male vs. female genomic coverage data (Methods and see **Supplementary Material 1** for details).

Overall, we identified between 139-246 contigs as putative Y-specific across the three different Y’s (using a cutoff of Log2(Male/Female) > 1 and >5x depth in males; **Table S8-S10, Figure 2**). In the Y_M_ assembly, several Y contigs have increased male coverage, indicating that they contain collapsed sequences. When we account for the inferred copy number of Y-specific contigs based on male coverage (**Table S8-10**), the assembled sizes are Y_S_=38.6Mb, Y_M_=43.9Mb and Y_L_ =59.5Mb. Importantly, the assembled sizes of the three Y chromosomes are in good agreement with our estimated sizes based on flow cytometry (**Table 1**), suggesting that we were able to recover a large fraction of the sequence of each Y chromosome in our assembly.

**Figure 2.**
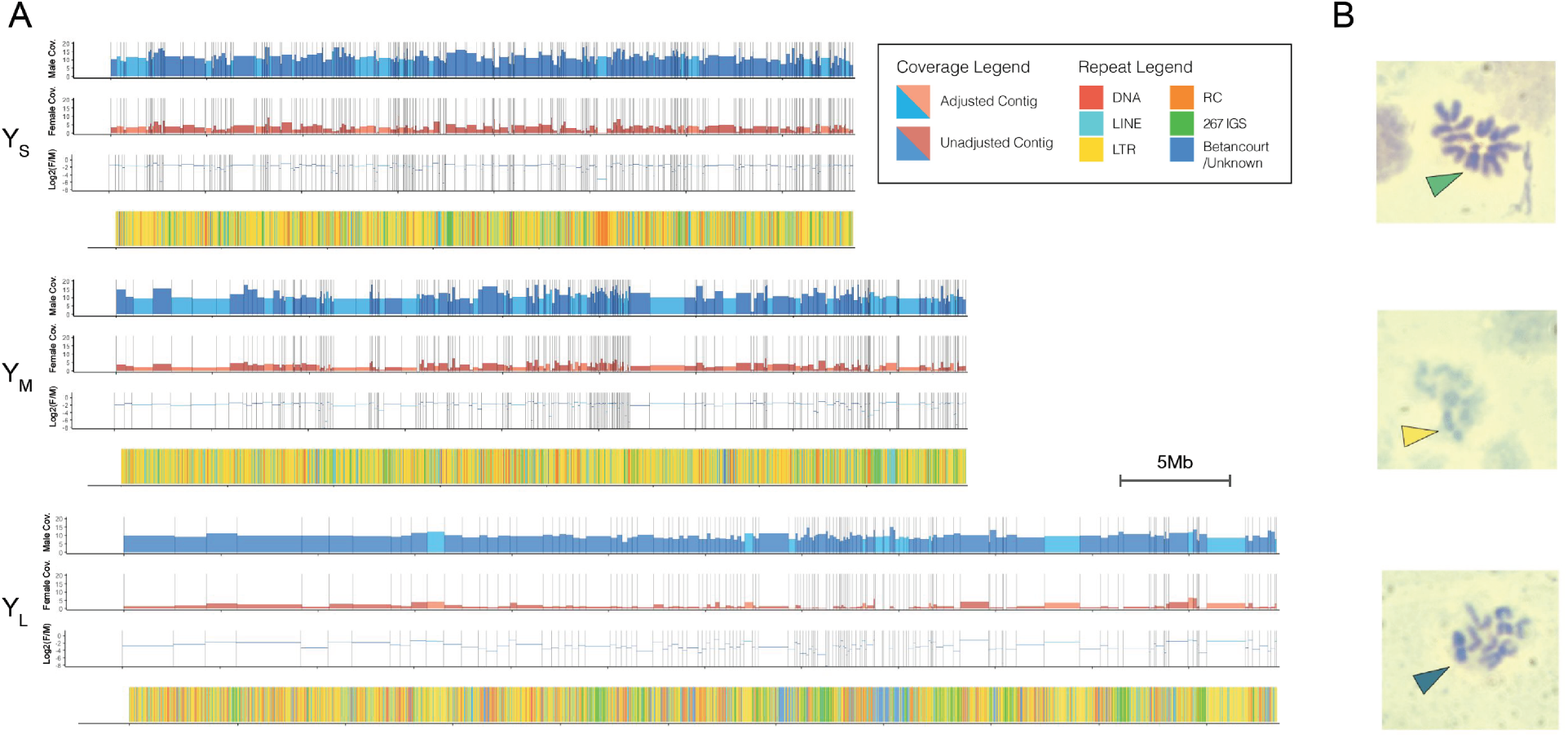
De novo assembly of three different sized Y chromosomes (YS, YM, YL). **A.** Male and female coverage tracks of Y-chromosome assemblies with Log2(Female/Male) coverage and repeat landscape shown beneath. One tick mark corresponds to 5Mb and collapsed contigs were adjusted in the plots. Landscapes of the most abundant repeats are shown separately in Figure S8 A-C. **B.** Top to bottom: karyotypes of YS, YM, YL males with the arrowhead denoting the Y chromosome.

**Table 1.** Summary of Y chromosome size estimations (in Mb) in *D. pseudoobscura* using different methods.

| Strain | Y<sub>S</sub> | Y<sub>M</sub> | Y<sub>L</sub> |
| --- | --- | --- | --- |
| Flow cytometry | 33 | 44 | 59 |
| Δ YS | -- | 11 | 26 |
| Assembly (total) | 38.6 | 43.9 | 59.5 |
| Δ YS | -- | 5.3 | 20.9 |
| dnaPipeTE | -- | 5.6 | 16.8 |
| TE library mapping | -- | 8.6 | 24 |
| Assembly Repeat Masked | 34.4 | 39.1 | 52.4 |
| Δ YS | -- | 4.7 | 18 |

We used BioNano to validate and further scaffold our assembly for Y_L_ (**Table S11, Supplementary Material 2**). Optical mapping produces restriction maps for long strands of DNA which can then be used to validate and place small contigs onto larger scaffolds. The BioNano hybrid assembly pipeline yielded 32 hybrid scaffolds, 6 of which contained the autosomes and X chromosome. The remaining 26 contained sequences from our Y_L_ Nanopore assembly. In total, 56 Nanopore contigs were fused into 17 hybrid scaffolds and the remaining 9 scaffolds contained sequences of single contigs. Overall, 63 of the 139 Y_L_ contigs are supported by optical reads, accounting for 41.7Mb of the 55Mb Y_L_ assembly, and only 15 contigs were cut. As expected, Nanopore contigs that were validated by optical mapping are significantly longer (mean length 662.5kb) vs. those not covered by optical reads (mean length 175.0kb). Thus, optical mappings validated 75.8% of our Y_L_ assembly (**Supplementary Material 2**).

### Molecular characterization of repeat content of Y chromosomes

The ancestral Y chromosome in *Drosophila* consists mostly of satellite and TE-derived DNA (26, 30). The Y chromosome of *D. pseudoobscura* is not homologous to the ancestral Y of *D. melanogaster* (31) but instead originated from an ordinary autosome about 15MY ago (32). Below, we use both assembly-free and assembly based methods to characterize the sequence content of the three different Y chromosomes at the molecular level.

As mentioned, cytogenetic methods suggest that Y_M_ and Y_L_ are 11Mb and 26Mb larger than Y_S_, with an absolute size of Y_S_=33Mb, Y_M_=44Mb and Y_L_ =59 Mb (**Table 1**). Most of this difference is likely driven by repeat accumulation (**Figure 1**). We used different metrics to quantify the repeat content of the different Y chromosomes, and estimate if differences in repeat content can account for the inferred size difference among Y chromosomes. In particular, we estimate the proportion of short reads assembling to TEs using dnapipeTE (33), and the proportion of short reads mapping to a curated TE library of *D. pseudoobscura* (34). Since the three different Y chromosomes are isogenic at the rest of the genome, these metrics can thus be used to estimate differences in the repeat content of the different Y’s.

As expected, we find that the Y replacement line containing Y_S_ has a larger fraction of single-copy reads (71.5%) followed by Y_M_ (69.9%) and Y_L_ (66.7%) (**Figure S4**). Using the difference in single-copy reads to estimate size differences between the Y’s suggests that Y_M_ and Y_L_ have an additional 5.6Mb and 16.8Mb of repeats in their genomes, respectively, in comparison to Y_S_ (**Table 1**). We also mapped raw Illumina reads to a TE reference library from *D. pseudoobscura* (34), to infer size differences in repeat content among Y chromosome, by normalizing coverage to the repeat libraries by autosomal coverage. Overall, we found that Y_S_ males contained about 85.3Mb of repetitive DNA, Y_M_ males contained 94.0Mb, and Y_L_ males 109.3Mb (**Table S12, Figure S5**). This suggests that Y_M_ and Y_L_ have an additional 8.6Mb and 24.0Mb repeats in their genomes, respectively, in comparison to Y_S_ (**Table 1**).

Lastly, heterochromatin-enriched genome assemblies recover overall similar differences in repeat content. We repeat-masked our three different Y chromosome assemblies using RepeatMasker version 4.1.0 with the *D. pseudoobscura* family repeat library (34). Indeed, the vast majority of Y-linked sequence is repetitive in all three assemblies, with total repeat-masked bases on Y_S_=34.4Mb (89%), Y_M_=39.1Mb (89%) and Y_L_ =52.4Mb (88%). Retroelements are by far the most abundant sequence class on each of the three different Y chromosomes (**Table 2**). We found that the TE families repeat-masked in our de novo assemblies corroborated the families we found in our Illumina mapping **(Table 1, 2; Table S13, Figure S6)**. TE abundance estimated from both Illumina mappings and de novo assemblies had a significant linear correlation for all Y chromosome variants (**Figure S6, p-value < 2e-16**).

**Table 2.**
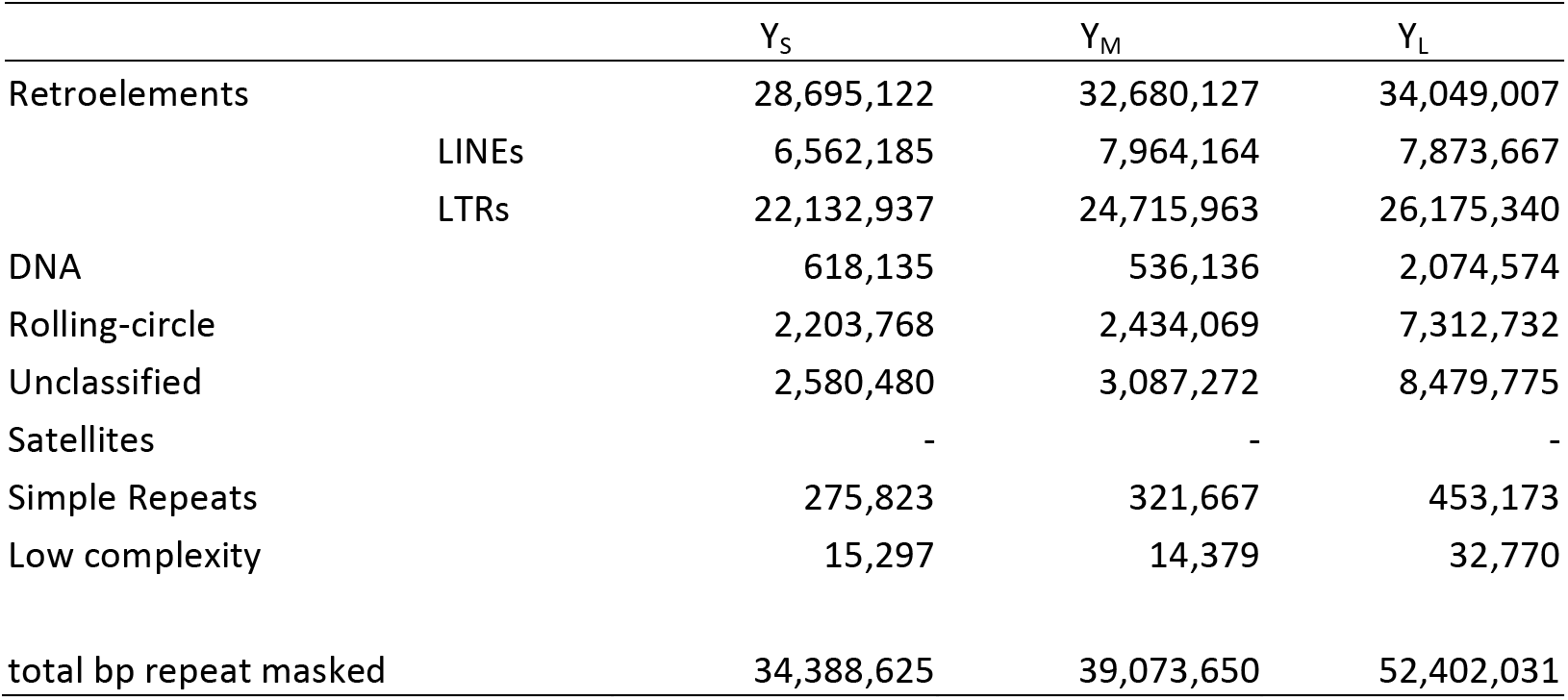
Summary of sequences repeat-masked in *D. pseudoobscura* Y chromosome assemblies.

Thus, these orthogonal approaches, using both cytogenetics and high-throughput sequencing, as well as assembly-based and assembly-free methods are largely in agreement about the different Y chromosome sizes. These results illustrate that our de novo Y assemblies are of high quality, and that the differences in Y size are largely driven by differences in repeat accumulation.

### Recent repeat accumulation drives size difference among Y chromosomes

The difference in sizes among Y chromosomes could be due to some repeats being mobilized on the Y, or due to structural mutations (such as duplications and deletions) resulting in size variation. Indeed, Dobzhansky hypothesized that smaller Y chromosomes originated via multiple deletions from the largest Y (19). To identify which repeats contribute to the observed size differences among Y chromosomes, we used both assembly-free and assembly-based methods. We mapped short reads to a repeat library, to infer different abundances of repeats on the different Y chromosomes (**Figure 3A, Table S12**), and also identified repeats on our assembled Y chromosomes (**Figure 3B, Table S13**). Intriguingly, we found that only a handful of repeats were responsible for the majority of additional DNA of Y_L_ and Y_M_ (**Table 3**). Interestingly, most of the TE families responsible for additional DNA of Y_L_ are not causing the size increase of Y_M_.

**Figure 3.**
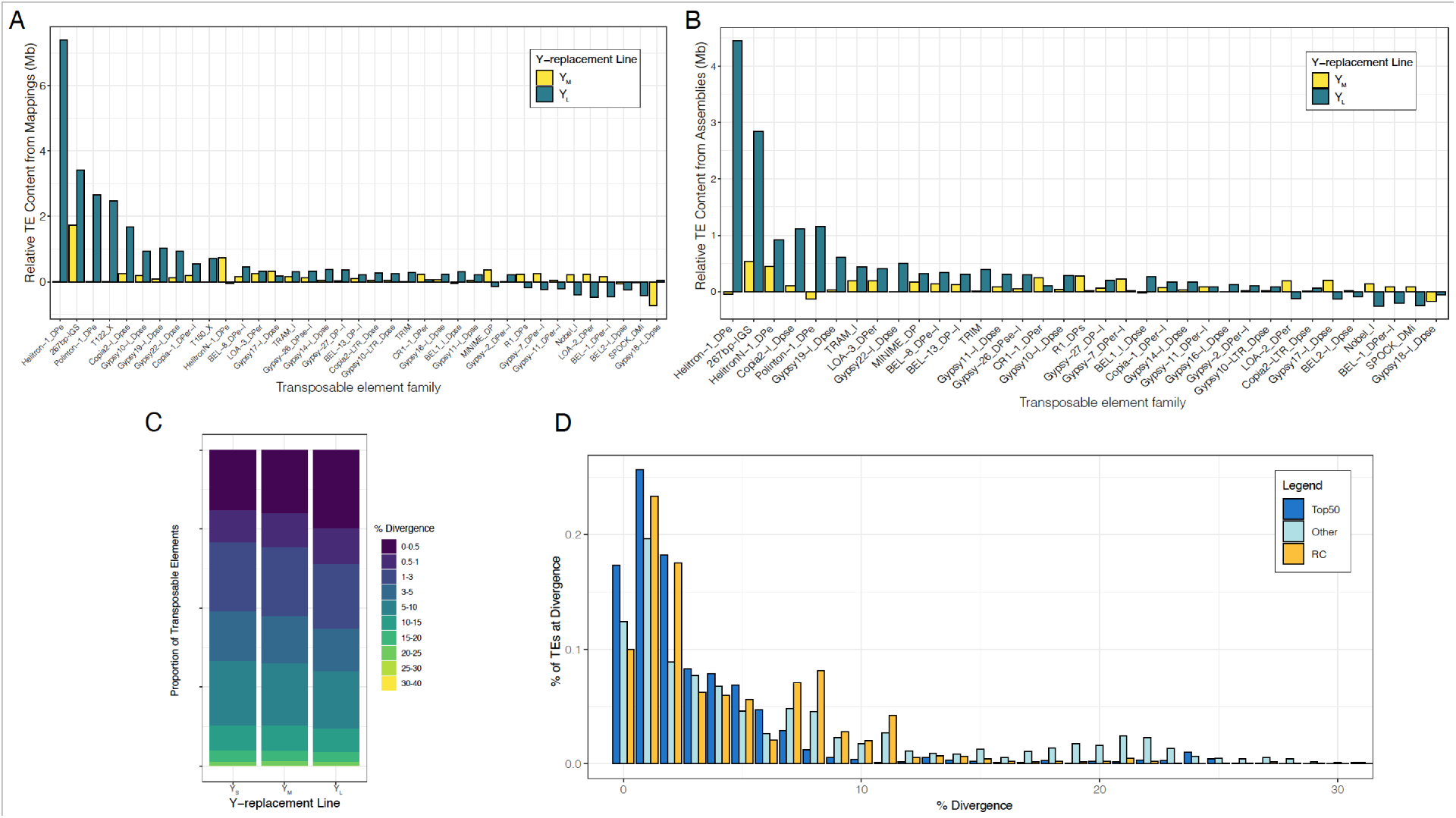
Repeat abundance and age suggests recent mobilization contributes to size differences among the Y’s. **A.** TE abundance in YM and YL relative to YS from paired-end mappings to the TE library. TEs where the absolute difference is over 200kb are shown. **B.** TE abundance in YM and YL chromosome assemblies relative to the YS assembly. TEs where the absolute difference is over 200kb are shown. **C.** Divergence in TE copies from whole-genome Illumina sequencing across the Y’s estimated by dnaPipeTE. **D.** Shown is the Kimura-based divergence in TE copies from the YL assembly based on the top 50 most abundant TEs (relative to YS), other TEs, and rolling-circle transposons. Percentages of TEs sum to 100% for each category.

**Table 3.**
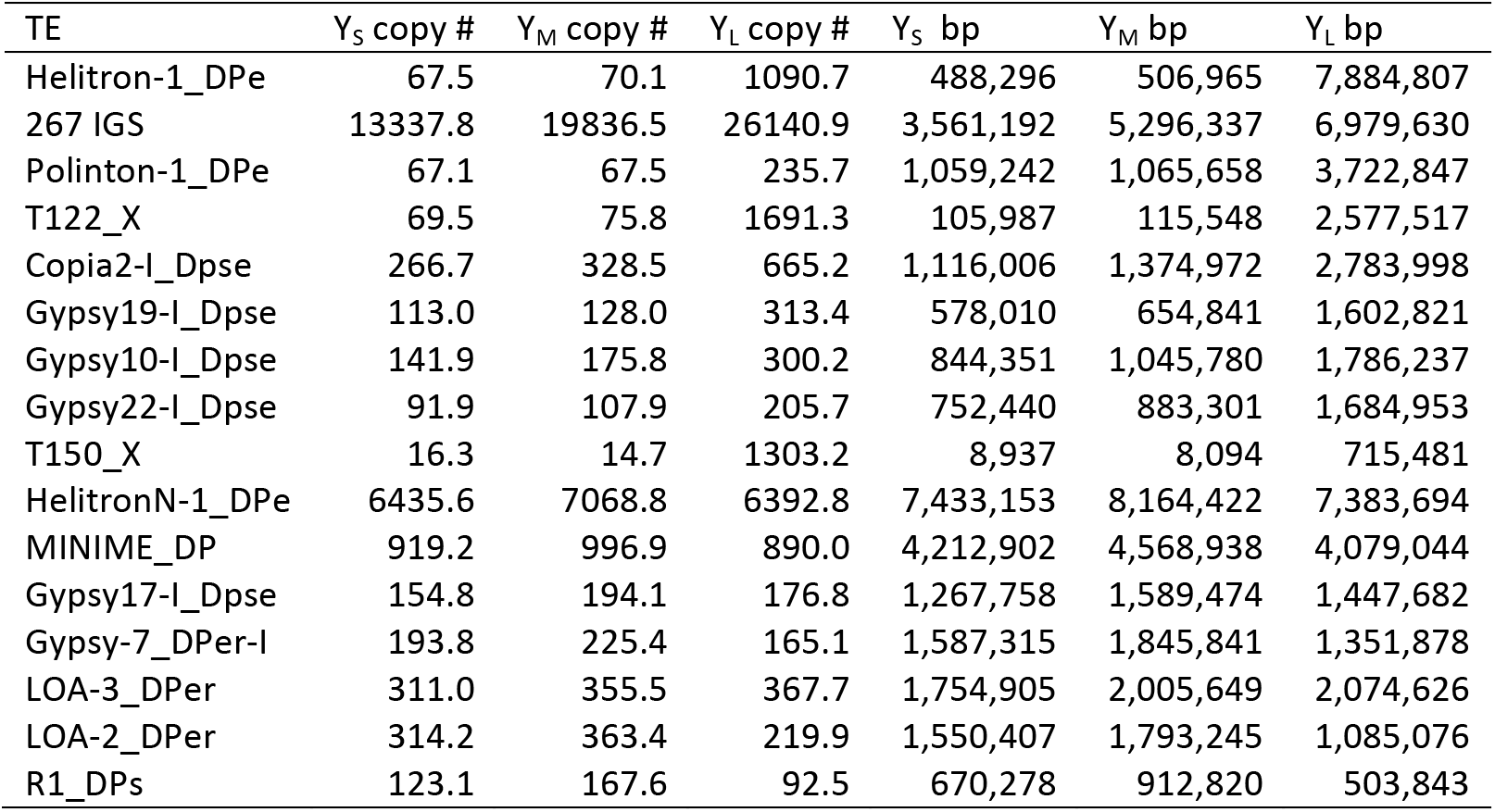
Size contribution (in bp) and inferred copy number of top repeats responsible for size differences of in *D. pseudoobscura* Y chromosome.

| TE | Y<sub>S</sub> copy # | Y<sub>M</sub> copy # | Y<sub>L</sub> copy # | Y<sub>S</sub> bp | Y<sub>M</sub> bp | Y<sub>L</sub> bp |
| --- | --- | --- | --- | --- | --- | --- |
| Helitron-1_DPe | 67.5 | 70.1 | 1090.7 | 488,296 | 506,965 | 7,884,807 |
| 267 IGS | 13337.8 | 19836.5 | 26140.9 | 3,561,192 | 5,296,337 | 6,979,630 |
| Polinton-1_DPe | 67.1 | 67.5 | 235.7 | 1,059,242 | 1,065,658 | 3,722,847 |
| T122_X | 69.5 | 75.8 | 1691.3 | 105,987 | 115,548 | 2,577,517 |
| Copia2-I_Dpse | 266.7 | 328.5 | 665.2 | 1,116,006 | 1,374,972 | 2,783,998 |
| Gypsy19-I_Dpse | 113.0 | 128.0 | 313.4 | 578,010 | 654,841 | 1,602,821 |
| Gypsy10-I_Dpse | 141.9 | 175.8 | 300.2 | 844,351 | 1,045,780 | 1,786,237 |
| Gypsy22-I_Dpse | 91.9 | 107.9 | 205.7 | 752,440 | 883,301 | 1,684,953 |
| T150_X | 16.3 | 14.7 | 1303.2 | 8,937 | 8,094 | 715,481 |
| HelitronN-1_DPe | 6435.6 | 7068.8 | 6392.8 | 7,433,153 | 8,164,422 | 7,383,694 |
| MINIME_DP | 919.2 | 996.9 | 890.0 | 4,212,902 | 4,568,938 | 4,079,044 |
| Gypsy17-I_Dpse | 154.8 | 194.1 | 176.8 | 1,267,758 | 1,589,474 | 1,447,682 |
| Gypsy-7_DPer-I | 193.8 | 225.4 | 165.1 | 1,587,315 | 1,845,841 | 1,351,878 |
| LOA-3_DPer | 311.0 | 355.5 | 367.7 | 1,754,905 | 2,005,649 | 2,074,626 |
| LOA-2_DPer | 314.2 | 363.4 | 219.9 | 1,550,407 | 1,793,245 | 1,085,076 |
| R1_DPs | 123.1 | 167.6 | 92.5 | 670,278 | 912,820 | 503,843 |

A few distinct repeats account for most of the size increase in the large Y chromosome (**Table 3**). In particular, we find that the Helitron-1_DPe transposon has notably higher copy numbers on Y_L_ compared to Y_M_ and Y_S_. Read coverage suggests that this TE (which is about 7.2kb in size) has roughly 70 copies in the Y_S_ and Y_M_ strain, but 1090 copies in Y_L_, resulting in an extra 7.4Mb of sequence on Y_L_ (**Figure 3A, Table 3; Figure S7**). A similar excess of this transposon is found in the assembled Y chromosomes: about 200kb and 150kb are repeat-masked on Y_S_ and Y_M_ for Helitron-1_DPe, but 4.7Mb on Y_L_ (**Table S13**). As expected for an active TE, we find copies of Helitron-1_DPe distributed across many different Y-linked scaffolds (**Figure S7, S8**). Helitron TEs replicate through a rolling-circle mechanism, which can result in the generation of tandem copies of the element (35). Indeed, genomic distribution patterns suggest that this element can occur in clusters of tandem arrays across the large Y chromosome (**Figure S8, S9**). BLAST searches using queries consisting of the last 50bp of the Helitron-1_DPe element fused to the first 50-bp yielded 50 hits on our Y_L_ assembly, and mate-pair violations using paired-end sequencing data further support the presence of partial copies of tandemly repeated elements (**Figure S9, Table S15**). Because many of the tandem copies of *Helitron-1_DPe* on Y_L_ are partial (**Figure S9**), our BLAST search yields a conservative estimate for the number of copies in tandem. We find 8 nearly full-length copies (using an 70% alignment cutoff) of Helitron-1_DPe on the assembled autosomes and X (and 3 copies on unmapped scaffolds). This is in contrast to the 289 near-full length copies we find in the Y_L_ assembly and the 10 and 14 copies in the Y_M_ and Y_S_ assemblies, respectively.

The repeat that is responsible for most of the size increase of Y_M_, and ranked second on Y_L_, is a 267-bp long satellite sequence that is homologous to the intergenic spacer region (IGS) of the rDNA cluster. In many Drosophila species, including *D. melanogaster*, the rDNA cluster is found on both the X and Y chromosome, and functions as a pairing site for the sex chromosomes during meiosis (36). A previous study has shown that the Y of *D. pseudoobscura* – which is not homologous to the Y of *D. melanogaster* – lacks functional rDNA clusters on the Y (37). Instead, *in situ* hybridization revealed four clusters bearing only the IGS of the rDNA repeats on the Y chromosome of the *D. pseudoobscura* strain investigated (37). We infer roughly 13,300 copies of the IGS satellite on Y_S_, 19,800 copies on Y_M_, and 26,100 copies on Y_L_, which adds about 1.7Mb of DNA to Y_M_, and 3.4 Mb of DNA on Y_L_ (**Table S12, Table S13**, **Figure 3A, 3B**). The IGS repeats are highly clustered in our assemblies, as expected for satellite sequences, and consistent with the *in situ* results (37).

The next most common repeat on Y_L_ is the Polinton-1_DPe repeat. Polintons are large DNA transposons (Polinton-1_DPe is 15.8kb) that encode several proteins necessary to replicate themselves (38). Read coverage analysis suggests that there are about 67 copies of this element on Y_S_ and Y_M_, but 236 copies on Y_L_, adding an additional 2.7Mb of sequence (**Table S12**). This element is distributed widely across different Y-linked scaffolds (**Fig S8**), and even read-coverage across the element suggests that many copies are full-length on the different Y chromosomes (**Fig S7**). The next most common element is the T122 repeat, a 1.5kb long uncharacterized element with roughly 70 copies on Y_S_ and Y_M_, but almost 1700 copies on Y_L_ (**Table S12**), adding an additional 2.5Mb of sequence. T122 repeats are often found in tandem clusters, mostly separated by HelitronN-1_DPe sequences, which may indicate co-amplification of T122_X by Helitron-initiated gene-capture. The next most common repeat is the Copia2-I_Dpse element (4.2kb in size) that has 270 and 330 copies on Y_S_ and Y_M_, respectively, but 670 on Y_L_, thereby adding 1.7Mb, and three different gypsy elements that are 2-3x more abundant on Y_L_ than Y_S_ and Y_M_, and each adding about another 1Mb of sequence to Y_L_. Thus, a total of only 8 repeats are responsible for over 20Mb of additional DNA on Y_L_ (i.e. >85% of the total size gain), with Helitron-1_DPe by far being the largest contributor to the size gain of Y_L_.

On the other hand, a much larger number of TEs contribute to the more modest size gain of Y_M_, with the top 8 expanded repeats contributing to <50% of size gain on Y_M_. As mentioned, the repeat leading to most of the size increase on Y_M_ is the IGS satellite (adding about 1.7Mb of DNA to Y_M_). The next most common one is the 1.1kb long HelitronN-1_DPe element and the related 4.6kb long MINIME_DP element (**Figure S10**), which have roughly about 6400 (HelitronN-1_DPe) and 900 (MINIME_DP) copies on both Y_S_ and Y_M_, but 7100 (HelitronN-1_DPe) and 1000 (MINIME_DP) copies on Y_M_ (thereby adding 730kb and 360kb of sequence to Y_M_). *Gypsy17-I_Dpse* adds another 320kb to Y_M_, and all other amplified TE families add less than 200kb sequence each to Y_M_ (**Table S12**).

We also used our assembled Y chromosome contigs and used Repeatmasker to annotate repeats and infer differences in abundance (**Figure 3B, Figure S11, Table S13, Table S14**). Because some contigs in our assemblies were collapsed, we calculated these differences by also considering the copy number of assembled contigs and their respective repeats. Similar to our assembly-free method, we found that Y_M_ and Y_L_ had 5.4 and 19.2Mb extra transposable elements, respectively, in comparison to Y_S_ (**Figure 3B**). Again, the same TE families (*Helitron*, 267bp IGS, *Copia, Polinton*) as found in the coverage-based approach are responsible for the size differences between Y chromosomes based on repeat-masking our assemblies. Thus, both methods suggest that only a small subset of repeats drive most of the observed size differences on Y_L_, and that there is overall little correspondence in the types of TEs that have accumulated on the different sized Y chromosomes.

Individual copies of recently active TEs are more similar to each other than TEs that are older, and the amount of sequence divergence of individual copies from the consensus can be used to infer the age of when a particular TE family proliferated (39). Consistent with the size increase on polymorphic Y’s being driven by the recent expansion of a few TEs, Y_M_ and especially Y_L_ contain younger TEs. We find that Y_M_ and Y_L_ have 4.9% and 23.3% more of their genome-wide TEs within 0-1% divergence from the consensus compared to Y_s_, respectively (**Figure 3C**). Further, the subset of TEs that are the main contributor to the larger sizes are younger than other TEs (**Figure 3D, Figure S12**), and amplified TEs cluster on phylogenetic trees (**Figure S13**). We further leveraged our assemblies to characterize divergence of TEs strictly located on the Y’s. As expected, we found many copies of TEs from different families, especially rolling circle elements (RC; to which the *Helitron* element belongs), were young in the large Y chromosome (**Figure 3D**). Thus, our de novo genome assemblies provide orthogonal confirmation that TEs have recently amplified on the large Y. Together, these results demonstrate that the large Y chromosome is the derived state in *D. pseudoobscura*, and DNA accumulation on these larger Y chromosomes is due to recent TE activity.

### Phenotypic consequences of repeat accumulation

While repetitive DNA is often viewed as a genetic wasteland, recent studies have found that differences in repeat content can have profound phenotypic consequences. In particular, it was shown that expression of 100s of genes differed in *D. melanogaster* males with polymorphic Y chromosomes (14, 15). Additionally, the presence and the number of Y chromosomes strongly influences genome-wide enrichment patterns of repressive chromatin modifications (40, 41), with additional Y chromosomes diminishing the heterochromatin enrichment at highly repetitive regions such as pericentromeres. Larger, more repeat- and gene-rich Y chromosomes were also linked to an up-regulation of transposable elements in males (42, 43). These results are generally interpreted as the Y chromosome acting as a heterochromatin sink that re-distributes repressive chromatin marks genome-wide (16). The integrity of heterochromatin deteriorates as individuals age (44), and the Y chromosome in *D. melanogaster* has been shown to contribute to heterochromatin loss and shorter longevity in males (41).

To test for phenotypic consequences of Y chromosome repeat content, we measured longevity of males that contained a small or large Y chromosome (Y_S_ and Y_L_) versus females, and assayed global transcriptome and chromatin profiles from Y_S_ and Y_L_ males. We collected H3K9me3 profiles from brains of young Y_S_ and Y_L_ males, in order to compare the global heterochromatin landscape in males with different Y chromosomes. The heterochromatin sink effect of the Y would predict that global H3K9me3 levels may be lower for repeats in males with a larger Y chromosome. **Figure 4A** shows that H3K9me3 enrichment is similar for TEs located on the X and autosomes in males containing differently sized Y chromosomes; however, TEs located on the Y chromosome show significantly lower H3K9me3 levels in Y_L_ males than Y_S_. Thus, the higher repeat content of Y_L_ may indeed dilute heterochromatin components in males with a large Y. Intriguingly, we find that H3K9me3 levels are generally much lower for TE copies located on the Y chromosome compared to autosomes and the X, irrespective of the Y variant (**Figure 4A, B; Figure S14, S15**). This is consistent with previous findings showing lower levels of heterochromatin enrichment for Y-linked TEs (42, 43), and repeat-rich Y chromosomes were found to be associated with the up-regulation of transposable elements in males (41–43).

**Figure 4.**
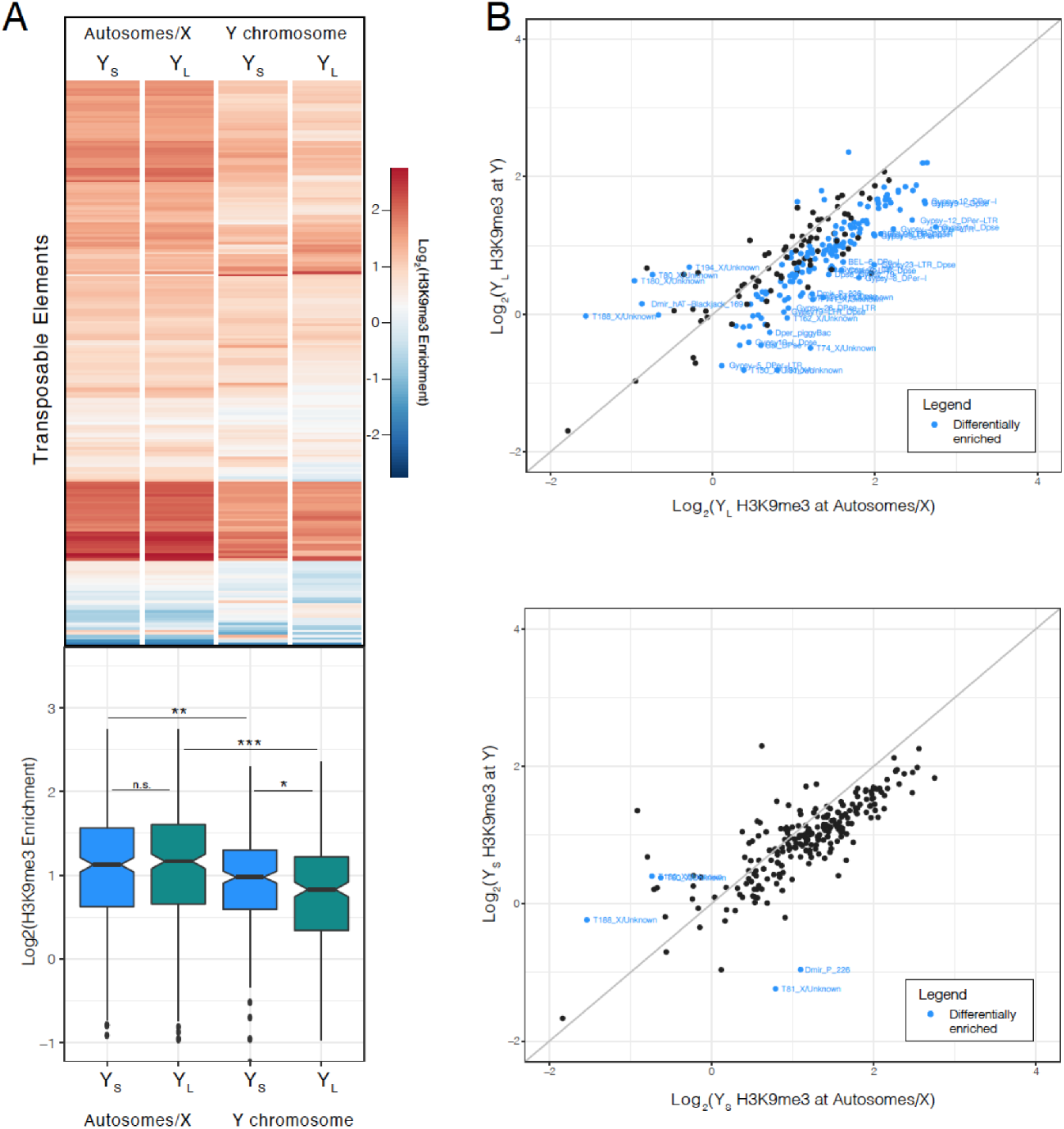
Decreased H3K9me3 enrichment on YL. **A.** Heatmap with corresponding boxplot showing H3K9me3 enrichment at TEs by chromosome (autosomes/X vs. Y chromosomes). Significance values calculated (* < 0.05, ** <0.01, *** < 1e-5, Wilcoxon-test). **B.** Scatterplot of H3K9me3 enrichment at TEs by chromosome group for YS and YL (p < 0.05, Two Sample t-test) The grey line indicates similar enrichment levels for both chromosome groups.

We obtained replicate expression data from heads of young and old Y_S_ and Y_L_ males to quantify TE expression between differently sized Y chromosomes, and during aging. While global expression across all TE families is not significantly different between Y-replacement males, irrespective of age **(Figure S16**), we find that a larger number of TE families are significantly up-regulated in Y_L_ males compared to Y_S_ (12 vs 0, at least 50% higher expression, Wald-test, p-value < 0.05; **Figure 5A**). Furthermore, half of these TEs are more abundant on Y_L_ than Y_S_, suggesting that the TEs that have accumulated on Y_L_ are responsible for increased TE expression (**Figure 5B**). Notably, this result is consistent irrespective of age, suggesting that differential regulation of TEs starts early in adult males (**Figure S17**).

**Figure 5.**
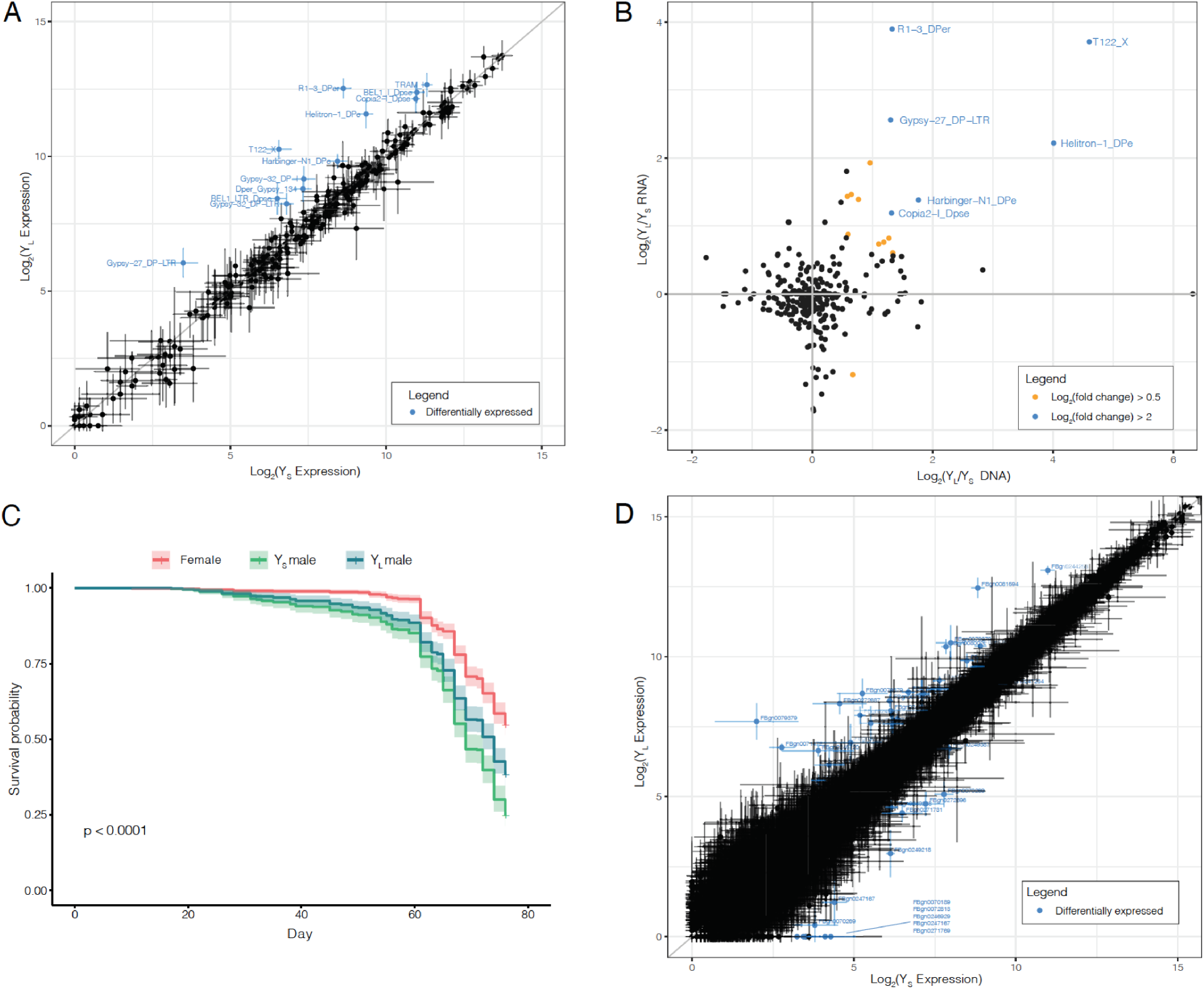
Differential transposon and gene regulation by Y chromosome. **A.** Differential TE expression between YS and YL males irrespective of age. Data represents the mean of 4 replicates with standard error bars (50% higher or lower expression, p < 0.05, Wald-test). **B.** Differential TE expression between YS and YL plotted against their differential TE abundance from Illumina mappings. **C.** Lifespan curves of YS and YL males and backcross females. **D.** Differential gene expression between YS and YL irrespective of age. Data represents the mean of 4 replicates with standard error bars (50% higher or lower expression, p < 0.05, Wald-test).

Interestingly, old Y_L_ males had slightly higher TE expression compared to young Y_L_ males but this was not true in Y_S_ males **(Figure S18**). Consistent with previous studies, we find that males live shorter than females (p-value < 0.0001). However, we find no significant difference in longevity between males containing the different sized Y chromosomes (**Figure 5C, Figure S19**); if anything, Y_L_ males live slightly longer than their counterpart Y_S_ males (median survival time 77 vs. 74 days, respectively). Thus, while males with a larger Y chromosome show slightly increased levels of TE expression, and more so in old males, these differences are relatively minor, and do not result in faster aging. Small differences in TE expression and lifespan are consistent with previously published results done in *D. melanogaster* Y replacement lines (13).

Previous studies in *D. melanogaster* have shown that different Y chromosomes can influence the expression of hundreds of genes located on the X and autosomes (15). We find that 930 genes on the autosomes and X exhibit differential expression between Y_S_ and Y_L_ Y-replacement lines, 20 of which are significant (>50% fold change, Wald-test, p-value < 0.05, **Figure 5D, Table S16**). Many differentially expressed genes are involved in the regulation of chromatin and transcription such as *aub, Ulp1*, and *mxc* (**Table S16**). Taken together, our results underline the impact Y chromosomes can have on genome-wide expression regulation.

### Repeated TE mobilization causes Y expansion in other Drosophila species

Our data suggest that modest changes in Y chromosome size are caused by copy number changes of a large number of repeats (Y_M_), but dramatic size differences are due to mobilization of a few elements (Y_L_). We used flow cytometry to study Y chromosome size polymorphism in five different Y-replacement lines in *D. affinis* (kindly provided by Rob Unckless), another member of the *obscura* species group and distant relative of *D. pseudoobscura*. We found three Y chromosomes that dramatically differed in size (**Figure 6**). Compared to the smallest Y chromosome of *D. affinis* (Y_S,*aff*_; strain YAF41), the medium-sized Y (Y_M,*aff*_; strain YAF159) has an additional 12.7Mb of DNA, and the largest Y chromosome (Y_L,*aff*_; strain YAF79) has an extra 19.0Mb of DNA (**Figure 6A**). We used Illumina sequencing of males with the differently sized Y chromosomes, and mapped reads to the same repeat library used for *D. pseudoobscura* to infer which repetitive elements are responsible for the size increase of the Y in *D. affinis*. Intriguingly, overall patterns of repeat accumulation on the large Y chromosomes are similar between *D. affinis* and *D. pseudoobscura*, with only a small number of TEs accounting for most of the size increase of the large Y chromosome (**Figure 6B, Table S17**). Most surprisingly, we find that the same TE (Helitron-1_DPe) that expanded on Y_L_ in *D. pseudoobscura* also is the main contributor to increased DNA content of Y_L,*aff*_ in *D. affinis*; 8 Mb of additional DNA (42%) on the large Y of *D. affinis* are due to additional copies of the Helitron-1_DPe element. The second most common repeat on Y_L,*aff*_ of *D. affinis* is Daff_Jockey_18 (3.4Mb), followed by Polinton-1_DPe (2.2Mb), the same element which also amplified on Y_L_ in *D. pseudoobscura*. Congruent with our findings in *D. pseudoobscura*, TEs on Y_M,*aff*_ and Y_L,*aff*_ also show low levels of sequence divergence that are commensurate with Y size (**Figure 6C**). Thus, recent amplification of a small number of TEs is also responsible for Y expansion in other species.

**Figure 6.**
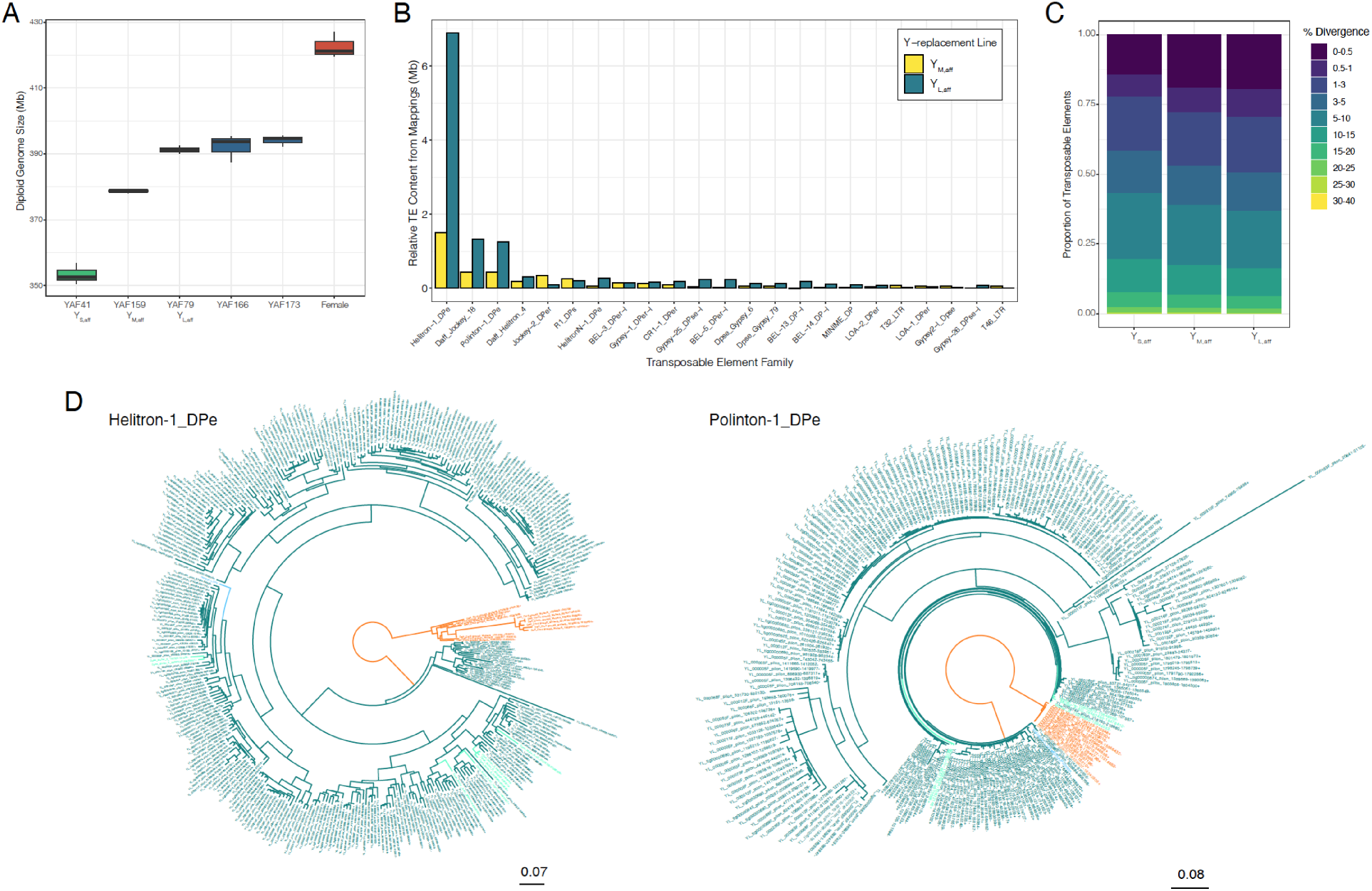
Y chromosome expansions in *D. affinis* by the same TEs as in *D. pseudoobscura*. **A.** Diploid genome size estimations from *D. affinis* Y-replacement lines. Rightmost boxplot (red) is the backcross female strain. **B.** TE abundance in YM, aff and YL, aff relative to YS, aff from paired-end mappings to the TE library, made in a similar manner as Figure 3B. TEs where the absolute difference is over 50kb are shown. **C.** Divergence in TE copies from whole-genome Illumina sequencing across the Y’s estimated by dnaPipeTE, made in a similar manner as Figure 3C. **D.** Phylogenetic trees of Helitron-1_DPe (left) and Polinton-1_DPe (right) found in *D. pseudoobscura* (green), *D. affinis* (orange), and *D.pseudoobscura* YL (teal). Repbase sequences are highlighted in blue.

TEs can transfer horizontally between species and this has previously been reported in the *D. pseudoobscura* group (34). To evaluate whether horizontal transfer between species could have contributed to the dramatic repeat accumulation on the large Y’s, we examined patterns of sequence evolution of the amplified Helitron and Polinton elements in *D. affinis* and *D. pseudoobscura*. We extracted Helitron-1_DPe and Polinton-1_DPe copies found on the autosomes/X of *D. pseudoobscura* and *D. affinis* and our Y_L_ assembly from *D. pseudoobscura*, and found that elements cluster by species (**Figure 6D**). Thus, this suggests that these elements evolved independently in the two species, as found in (34). Since we lack a high-quality assembly of the large Y chromosome in *D. affinis* we cannot reconstruct the sequence of individual Y-linked TEs in this species, but our coverage analysis implies that a large majority of the Helitron and Polinton reads are derived from Y_L_ and Y_L,*aff*_. We therefore constructed consensus sequences for these two TEs for both species and mapped genomic reads from *D. affinis* and *D. pseudoobscura* males with the large Y’s to the consensus sequences to estimate the rate of species-specific mapping. As expected if TEs amplified independently in the two species, most Y_L_ reads map to the *D. pseudoobscura* consensus and most Y_L,*aff*_ reads map to the *D. affinis* consensus (**Table 4**). Thus, we find no signatures of horizontal transfer at these elements, which suggests that rampant amplification of autonomous TEs occurred independently on the two different species’ Y chromosomes.

**Table 4.** Y_L_ and Y_L_,_aff_ Helitron/Polinton mappings to the *D. pseudoobscura* and *D. affinis* consensus sequences.

| Consensus sequence | Helitron-1_DPe derived reads |  | Polinton-1_DPe derived reads |  |
| --- | --- | --- | --- | --- |
|  | *D. pseudoobscura* | *D. affinis* | *D. pseudoobscura* | *D. affinis* |
| *D. pseudoobscura* | 562,655 | 2,126 | 265,181 | 10,963 |
| *D. affinis* | 5,670 | 349,819 | 399 | 98,907 |

## Conclusions

In a series of papers, Dobzhansky cataloged variation in size and shape of the *D. pseudoobscura* Y chromosome across its geographic range (19). However, the molecular basis or the mechanism of the origin of variation of Y size was unknown, and Dobzhansky speculated that the polymorphic Y’s may have been derived by losses of sections from the largest Y chromosome (19). Here, we use a combination of classical cytogenetic methods and long-read sequencing to identify the molecular basis of Y size variation. We show that in contrast to Dobzhansky’s hypothesis, most of the size differences between morphologically distinct Y chromosomes are due to the recent expansion of a few TEs on the largest Y chromosome. Thus, rather than being the ancestral form, this indicates that the large Y chromosome is instead derived from a smaller ancestral Y. Most intriguingly, we find that the same TEs families can contribute to Y expansion independently in different species of Drosophila. Dobzhansky also tried to find a correlation between the type of Y chromosome present in a given strain and other properties of that strain, including hybrid sterility (19, 45, 46), but concluded that the morphological and physiological characteristics of a particular male are not affected by the type of its Y chromosome. By homogenizing the genetic background of the different Y types, we were able to detect subtle differences among Y chromosomes. We show that overall heterochromatin enrichment is reduced on the Y of males containing a larger Y chromosome, and expression of a subset of TEs, and especially those that have accumulated on Y_L_, is increased. Additionally, expression of a few dozen of X-linked and autosomal genes is influenced by Y-type. Thus, Y variation can potentially influence male-related fitness traits.

Morphological variation in Y chromosome size and shape has been identified in many organisms, including humans (47), but has not yet been studied at the DNA sequence level. Our study demonstrates that novel sequencing approaches allow the assembly of even highly repetitive Y chromosomes, and presents novel insights into the repetitive landscapes of variable Y chromosomes.

## Acknowledgements

We thank Emily Chong, Amy Weixiang Wang, Kamalakar Chatla and Reema Aldaimalani for help with fly work and technical assistance, and members of the Bachtrog lab, especially Kevin HC Wei and Ryan Bracewell, for help with data analysis. We thank Carl Hjelman for advice on flow cytometry. We thank Spencer Johnston, Stephen Schaeffer, Nitin Phadnis, Ryan Bracewell, Rob Unckless and the Cornell Stock Center for providing fly strains.

## Materials & Methods

### Y-replacement line backcrosses

A total of 26 different *D. pseudoobscura* males were backcrossed to the sequenced *D. pseudoobscura* stock (MVZ 25, courtesy of Stephen Schaeffer). The list of fly strains used for Y-replacement lines is given in **Table S1**, and **Figure S1** shows the geographic origin of different lines. We backcrossed males to virgin females for 9 generations after which the Y-replacement lines were maintained on standard molasses culture. We verified homozygosity on the autosomes and X chromosome by mapping Illumina data to the masked genome, calling sites with Samtools mpileup, and then dividing the number of sites that are homozygous across all strains by the total number of variant sites (**Table S18**).

### Y chromosome karyotyping

We dissected brains from third instar larvae in 0.9% NaCl. First, we incubated brains in 0.9% NaCl with 2 drops of 0.1% colchicine for 10 min at room temperature, followed by another 10 min incubation in 0.1M KCl solution. Then, we fixed brains in 3:1 methanol and acetic acid for 1 hour. We moved the fixed tissues to 50μL of 60% acetic acid solution and vigorously disassociated tissues via pipetting for 1 minute. We dispensed the solution onto pre-heated 55°C microscope slides and stained them in 7% Giemsa solution (1x PBS pH 6.5) for 40 min. Slides were imaged on a Leica DM5000B microscope with SPOT imaging software. We then measured chromosome lengths with KaryoType (48).

### Flow cytometry

We estimated the sizes of Y chromosomes using flow cytometry as described (49). Briefly, we dissected and snap-froze heads from flies aged 3-7 days old. For an internal standard, we used heads from a *D. virilis* strain provided by Spencer Johnston (1C = 328Mb). To prepare samples, we dounce homogenized one *D. pseudoobscura* head with one *D. virilis* female head in a 2mL Kontes dounce homogenizer with 1mL Galbraith Buffer (45mM MgCl2, 30mM sodium citrate, 20mM MOPS, 0.1% Triton X-100 (v/v), pH 7.2) and filtered the nuclei-containing solution through a 100 μM cell strainer. We then added 25μL propidium iodide (1mg/mL) to stain nuclei for 30 min at 4°C and ran the stained nuclei on a BD LSR II cytometer at 30-60 events/sec.

We exported raw fluorescence data on FlowJo (v.10.1) and ran a custom R script to calculate the genome sizes. To calculate the genome size of the unknown Y-replacement line, we assumed fluorescent linearity and used the following equation:

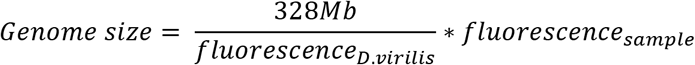

To estimate %AT content relative to *D. virilis*, we followed (49) and used 1μL of DAPI (1mg/1mL) in place of propidium iodide. We calculated %AT assuming linearity.

We estimated the proportion of under-replicated DNA in polytene tissue following the protocol described (22). Briefly, we dissected thoracic tissue from flies and followed the protocol described above without the *D. virilis* standard. We ran the same protocol with head tissue as a positive control. We used a custom R script to calculate the 4C cell fluorescence in relation to 2C fluorescence. All flow cytometry work was done at the UC Berkeley Cancer Research Lab Flow Cytometry Facility.

### Lifespan assays

We conducted lifespan assays following (50) with the following rearing conditions: 18°C, 60% relative humidity, 12h light, and standard Bloomington food. Briefly, we collected synchronized embryos on a molasses plate with yeast paste for 16-20 hours. We washed embryos with 1x PBS pH 7.2 three times and dispensed 10μL of embryos per culture vial by pipette. To obtain synchronized adults, we collected emerging adults over a 3 day window and aged the adults for 3 days to mate and copulate. Male and female flies were separated into separate vials, placing 30 flies per vial. We moved flies to new vials every 2-3 days without CO_2_ and recorded the number of deaths. Flies that escaped were censored. In total, 1196 sequenced strain female flies, 722 sequenced strain Y_S_ male flies, and 987 Y_L_ male flies were tracked for the lifespan assays recorded here.

### Tissue collection

Tissue for gDNA-Seq was collected by freezing male flies and popping off heads via vortexing. To collect tissue for the RNA-Seq experiment, we censored the entire experiment once it reached 50% survivorship for one line. We collected head tissue for RNA-Seq in a similar manner to the gDNA-Seq. For the ChIP-Seq experiment, we anesthetized young (5-7 day old) flies using CO_2_, dissected the brains in 1x PBS pH 7.2, and immediately froze 2 brains per sample using dry ice.

### DNA, RNA, and ChIP library preparation

We extracted DNA from pooled heads of each Y-replacement male using the DNAEasy Qiagen Extraction kit. We then prepared gDNA libraries using the Illumina TruSeq Nano DNA library kit (#20015965). For the cDNA libraries, we first extracted RNA from 20 pooled heads for Y-replacement line males using TRIzol (Invitrogen #15596026). We then made libraries using the Illumina TruSeq Stranded Total RNA Ribo-Zero Gold kit (Illumina RS-122-2201).

For the ChIP-Seq libraries, we started with brain tissue and prepared ChIP pull-downs as described in (43). Briefly, we modified the native and ultra-low input ChIP protocol (51) for double-brain ChIP-Seq. First, we added 40μL EZ nuclei lysis buffer to frozen tissues and homogenized them with a pestle grinder on ice. We spun cells down at 1000 g for approximately 10 min and decanted 20μL of the supernatant. We froze the cell pellet in the lysis buffer at −80°C. We then followed the 100,000 cell count digestion protocol from (51) with a few modifications. For each *D. pseudoobscura* sample and *D. melanogaster* spike-in sample, we added 4.4μL of 1% DOC and 1% Triton-X100 to resuspended cells and mixed thoroughly. Briefly, we digested each sample at 37°C for 04:58 (mm:ss) with MNase and quenched the reaction with 4.9μL of 100mM EDTA pH 8.0. We then added the fragmented spike-in samples to each *D. pseudoobscura* sample to account for approximately 20% of the final pooled sample volume (i.e. final sample consisted of 80% *D. pseudoobscura*, 20% *D. melanogaster*). We reserved 10% of the pooled fragmented sample for the input (fragment control) and used the remaining 90% to perform the chromatin pull-down (ChIP sample, target antibody: H3K9me3 polyclonal classic Diagenode C15410056) according to the 100,000 cell specifications. We used the Rubicon Genomics ThruPlex kit to prepare ChIP-Seq libraries for sequencing with 10 PCR amplification cycles for the input samples and 12 cycles for the ChIP pull-down samples. We sequenced libraries on the Illumina HiSeq 4000 Platform at the Vincent J. Coates Genomics Sequencing Center (Berkeley, CA).

### Repeat abundance estimations by reference library

We used the *D. obscura* group consensus TE library from (34) and included a Y chromosome-specific 267bp repeat sequence which was later confirmed as the 267bp intergenic spacer (37). We masked the *D. pseudoobscura* or *D.affinis* genome using this repeat library with RepeatMasker (-no_is -no_low -norna, v4.1.0). Then, we mapped reads with bwa mem (60) to the repeat-masked genome and the TE library. In this way, all reads derived from TEs will map to the consensus library to decrease reference bias for the small (reference) Y. We used bedtools coverage to calculate coverage per TE and normalized coverage by the median autosomal coverage. The autosomes were confirmed to have approximately 2x coverage (i.e. diploid genome).

We summed the copy numbers of all TEs multiplied by their respective lengths to estimate the difference in TE bp contribution between Y-replacement lines. Note that this sum includes TEs derived from autosomes, however, these contributions should be negligible because the strains have isogenic backgrounds.

### Repeat content estimation and divergence by dnaPipeTE

We ran dnaPipeTE on the first strands of samples with the following parameters: -coverage 0.5 -genome-size 175000000 -sample-number 2. Note that we used genome size of 175 Mb even though the strains are of different sizes, which will yield a more conservative repeat % result between the strains because of varying genome sizes. We used the % difference in single-copy elements to estimate the Mb difference between Y’s. We also used the estimated divergence to identify recently active TEs on a genome-wide basis.

### Nanopore sequencing and Y-chromosome de novo assembly

For the reference strain male (Y_S_), we used reads from (21). For Y_M_ males, we used one Nanopore MinIOn flow cell (v9.4.1 RevD) with the ligation sequencing kit (SQK-LSK109) and obtained 540,115 total filtered reads with a mean read length of 15,972bp. We sequenced Y_L_ males using 3 Nanopore MinIOn flowcells (v9.4) with the ligation sequencing kit (SQK-LSK108 and SQK-LSK109). We combined the total output from these three runs and obtained 482,379 filtered reads with a mean read length of 19,650bp.

We followed a heterochromatin-enriched assembly approach from (26). Briefly, we mapped reads to the *D. pseudoobscura* assembly (21) using minimap2 (v2.1) with default parameters. Then we extracted reads that did not map to putative euchromatin in the *D. pseudoobscura* assembly (21) to make heterochromatin-only assemblies using Canu (v1.6) (27) and Falcon (v1.4.2) (28) independent of each other.

We then removed contigs containing bacterial DNA using BlastN against the NCBI database and then used QuickMerge to merge contigs between Canu- and Falcon-based assemblies. We settled on assemblies that had a high N50 and generally less contigs, all of which resulted from merges between assemblies from a Canu to Falcon direction with assemblies from a Falcon to Canu direction. To identify Y contigs, we mapped Illumina male and female reads to the corresponding heterochromatin-based assemblies with bwa mem and obtained genomic coverages with bedtools coverage. Y chromosomes can contain repeats found elsewhere in the genome so we used differences in female and male coverages to identify male-bias/Y-linked sequences. Contigs whose Log2(Female/Male) coverage was below −1 and whose overall male coverage was at least 5x were considered Y contigs. We then appended these Y contigs to the assembled autosomes and X scaffolds to generate full male genomes. Using our coverage analysis, we also split contigs whose coverage was partially male-biased for the medium and small Y assemblies. Additional information can be found in **Supplementary Material 1**.

To calculate the estimated amount of DNA and repeats on each Y chromosome variant, we normalized the coverage of the Y contigs by the coverage of the autosomes. We used this normalized coverage as the corresponding copy number (estimated abundance) of each Y contig with which we used to adjust the estimated total DNA length and total repeat abundance (RepeatMasker v4.1.0).

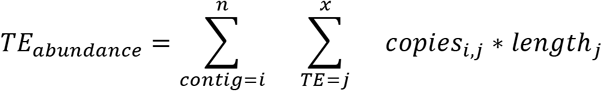

### BioNano molecules and hybrid assembly

We used optical mapping (BioNano Genomics) to stitch Y contigs from the large Y chromosome together. Briefly, DNA was extracted from 45mg of frozen male larvae using the Bionano Prep Animal Tissue DNA Isolation Kit (Product #80002) with modifications from a protocol by Susan Brown at Kansas State University. Then, DNA was homogenized and embedded in agarose plugs, treated with Proteinase K and RNAse A, and isolated from agarose and cleaned via drop dialysis. DNA was then left at room temperature for several days to homogenize prior to quantitation. 750 ng of DNA was labeled using the DLS Labeling Kit (Product #80005) following the Bionano Prep Direct Label and Stain (DLS) Protocol (Document 30206).

Labeled DNA was run through the Bionano Saphyr Platform. These reads were then assembled into full optical maps and then allowed for stitching of Y chromosome sequences. In total, 334,498 molecules were generated with an N50 of 223.875 kb and this was downsampled to proceed with the hybrid assembly pipeline (i.e. 189,039 molecules and N50 of 273.571 kb). The Bionano hybrid assembly contained 32 hybrid scaffolds, 6 belonging to the autosomes and X chromosome and the remaining 26 to Y_L_ chromosome. We confirmed the 6 scaffolds were from the autosomes and the X chromosome with NUCmer (v3.1) alignments to the *D. pseudoobscura* genome. Additional information can be found in **Supplementary Material 2**. The BioNano preparation and hybrid assembly were completed at the UC Davis DNA Technologies Core.

### Y-linked TE divergence estimates

To obtain Kimura divergence estimates for each TE family and their copies on the Y’s, we used parseRM_getLandscape.pl from (52) with our RepeatMasked Y chromosome assemblies.

### Tandemly repeated transposable element analysis

We adapted an approach from (53) to infer if transposons were tandemly repeated by identifying violations in paired-end alignments. Paired-end reads were first aligned with bwa mem (60) in single-end mode to the masked *D. pseudoobscura* genome and the TE library. We then obtained read pairs where both forward and reverse reads aligned to the same TE and then plotted their alignment coordinates.

To infer the number of tandemly repeated *Helitron-1_DPe* transposons in the Y_L_ assembly, we made artificial queries of full head-to-tail sequences as in (35). We used BLASTN (v2.6.0+) to blast this sequence to the Y assemblies and identified tandem junctions based on matches with > 80% aligned.

### Transposable element phylogenies

We used BLASTN (2.6.0+) to extract transposon sequences from our Y chromosome assemblies. We kept alignments of at least 70% length with the exception of Gypsy22-I_Dpse, Copia-1_DPer-I, Polinton-1_DPe, and T150_X where we kept alignments mapping to regions of elevated coverage. We used clustalo (v1.2.4) using default parameters to generate multiple sequence alignments and then RAxML (v8.2.12) using the following parameters: -f a -x 1255 -p 555 -# 100 -m GTRGAMMA. Phylogenies were visualized using ggtree (54) and FigTree (55).

### Gene expression analysis

Briefly, we mapped RNA-seq reads for each replicate to a repository of ribosomal DNA scaffolds from NCBI and removed all reads that mapped to this scaffold. Differences in rRNA abundance in sequenced samples are likely to be technical artifacts following RNA library preparation. We then mapped the remaining reads to the *D. pseudoobscura* genome and to the repeat library separately using HISAT2 (56) with default parameters. We then used Subread FeatureCounts (57) to calculate gene counts and repeat counts and used DESeq2 (58) to normalize the libraries and perform differential expression analysis between male/female flies of different ages.

### ChIP enrichment analysis

We followed the approach from (43). We took ChIP mappings to the entire genome and called enrichment at repetitive elements through TECounts (part of TETranscripts) (59). To call for repeat enrichment on a per chromosome basis, we split mappings by chromosome (autosome/X chromosome and Y chromosome) and ran TECounts separately for each split mapping. We then normalized counts at repeats in both the ChIP and input by mean autosomal coverage and calculated ChIP enrichment as ChIP/Input.

### Transposon cross-mapping analysis

We identified autosome/X Helitron and Polinton elements in both the *D. pseudoobscura* and *D. affinis* genomes using BLASTN, keeping alignments with at least 70% length. We generated consensus sequences using EMBOSS cons with default parameters (https://www.ebi.ac.uk/Tools/msa/emboss_cons, accessed April 22, 2021). For Y_L_ and Y_L,*aff*_ samples, we extracted reads mapped to Helitron-1_DPe/Polinton-1_DPe. Then, we used bwa mem (60) to map reads to both consensus copies of the corresponding transposon simultaneously.

## Supplementary Figure Legends

**Figure S1.** Geographic location of sampled *D. pseudoobscura* Y chromosomes.

**Figure S2.** Backcrossing scheme to generate Y-replacement lines.

**Figure S3.** Representative chromosome squashes of Y-replacement line males.

**Figure S4.** Reference-free based TE and single-copy (euchromatin) estimates from DNAPipeTE.

**Figure S5.** Total TE abundance derived from Illumina mapping. Data underlying figures is contained in Table S11. Note these are absolute amounts compared to the relative amounts which are plotted in Figure 3A.

**Figure S6.** Correlation between Illumina mapping and de novo assembly TE abundances. Each data point represents a TE family and its respective abundance calculated from Illumina mappings and repeat-masked Y chromosome assemblies. Data underlying these figures are contained in Table S11 and Table S12.

**Figure S7.** Coverage of top 10 most abundant transposons in Y_L_ from Illumina mappings for (A) Y_L_ and (B) Y_M_.

**Figure S8 (A-C).** TE distribution on Y-linked scaffolds for (A) Y_L_, (B) Y_M_, and (C) Y_S_. One unit corresponds to a 1-5kb window. Note that assemblies are not not adjusted for contig coverage.

**Figure S9.** Inferring tandemly repeated TEs using violations of forward and reverse mapping coordinates. TEs that have head-to-tail tandems will have clusters of reads in the bottom right corner. For each TE, reads are downsampled to the smallest Y chromosome sample with the exception of T150_X where Y_M_ has no reads.

**Figure S10.** Alignment of HelitronN-1_DPe (X-axis) to MINIME_DP (Y-axis) using nucmer.

**Figure S11.** Repeat composition on a per chromosome basis.

**Figure S12.** Divergence of abundant/non-abundant TEs on Y_M_ relative Y_S_.

**Figure S13.** Unrooted phylogenies of TE copies found in Y_S_, Y_M_, Y_L_ assemblies. D. miranda copies are identified for Copia2-I_Dpse, T122_X, and Helitron-1_DPe.

**Figure S14.** H3K9me3 enrichment for TEs with 50% more copy number in either Y_S_ or Y_L_.

**Figure S15.** Log2(H3K9me3 TE_Y_ / H3K9me3 TE_Auto/X_) for all Y_S_ and Y_L_ ChIP replicates.

**Figure S16.** TE expression by Y-replacement line and age, sorted by TE abundance. (A) All TEs with Log2(expression) > 1. (B) All TEs with Log2(expression) > 1 and 50% more TE abundance in either Y_S_ or Y_L_.

**Figure S17.** TE abundance differences vs TE expression differences in Y-replacement line males by age. Each data point represents a TE with its associated difference in abundance/expression between Y_S_ and Y_L_. Plots were made from the same data that underlies Figure 4C.

**Figure S18.** Age-associated TE expression differences within Y-replacement line males. Left: Y_S_ males, right: Y_L_ males. Blue data indicates at least 50% significant differential expression (Wald-test, p-value < 0.05).

**Figure S19.** Y-replacement line aging trials recorded separately. All aging trials are shown with females from both Y-replacement line males plotted as individual tracks.

## Supplementary Table Legends

**Table S1.** Starting wild-type strain information.

**Table S2.** Inferred Y chromosome size by karyotype measurements.

**Table S3.** Inferred Y chromosome morphology by chromosome arm ratios (data underlying Figure 1B).

**Table S4.** Y-replacement line male genome size estimates and morphology (data underlying Figure 1C).

**Table S5.** Heterochromatin estimates of Y-replacement lines (data underlying Figure 1D).

**Table S6.** Summary of genomic sequencing data and statistics.

**Table S7.** Summary statistics of Y chromosome assemblies. Note that the assemblies are not adjusted for the copy number of each contig.

**Table S8.** Contigs, their length, bp repeats masked, and M and F and inferred coverage ratio in Y_S_ assembly.

**Table S9.** Contigs, their length, bp repeats masked, and M and F and inferred coverage ratio in Y_M_ assembly.

**Table S10.** Contigs, their length, bp repeats masked, and M and F and inferred coverage ratio in Y_L_ assembly.

**Table S11.** BioNano hybrid scaffolds for Y_L_ assembly (lengths in Mb). Additional information can be found in Supplementary Material 2.

**Table S12.** Transposable element base pair contribution by Illumina data. Numbers represent whole-genome estimates and any differences will be due to the Y. Data underlies Figure 3A.

**Table S13.** Transposable element base pair contribution from Y chromosome assemblies (i.e. exclusively Y-linked contigs). Estimates are derived from RepeatMasker. Data underlies Figure 3B.

**Table S14.** Transposable element base pair contribution from female genome assembly (i.e. exclusively autosomes and X chromosome contigs). Estimates are derived from RepeatMasker. Data underlies part of Figure S11.

**Table S15.** Head-to-tail chimeric junction locations on Y assemblies.

**Table S16.** Differentially expressed genes between Y_S_ and Y_L_ strains. Up-regulation means higher expression in Y_L_ than Y_S_. Duplicated genes are due to multiple gene copies on the chromosomes in the *D. pseudoobscura assembly*.

**Table S17.***D. affinis* transposable element base pair contribution by Illumina data. Numbers represent whole-genome estimates and any differences will be due to the Y. Data underlies Figure 6A.

**Table S18.** Homozygosity on the autosomes/X chromosome for Y-replacement lines.

## Notes

### Competing Interest Statement

The authors have declared no competing interest.

## References

1. Bachtrog D (2020) The Y Chromosome as a Battleground for Intragenomic Conflict. Trends Genet 36 (7):510–522.

2. Bull JJ (1983) Evolution of sex determining mechanisms (Benjamin/Cummings, Menlo Park) doi:10.1093/molbev/msn020”,”keywords“:[”molecular.

3. Bachtrog D, et al. (2014) Sex determination: why so many ways of doing it? PLoS Biol 12(7):e1001899.

4. Charlesworth B, Charlesworth D (2000) The degeneration of Y chromosomes. Philos Trans R Soc Lond, B, Biol Sci 355(1403):1563–1572.

5. Bachtrog D (2013) Y-chromosome evolution: emerging insights into processes of Y-chromosome degeneration. Nat Rev Genet 14(2):113–124.

6. Charlesworth B (1978) Model for evolution of Y chromosomes and dosage compensation. Proc Natl Acad Sci USA 75(11):5618–5622.

7. Rice WR (1984) Sex chromosomes and the evolution of sexual dimorphism. Evolution 38(4):735–742.

8. Charlesworth B, Sniegowski P, Stephan W (1994) The evolutionary dynamics of repetitive DNA in eukaryotes. Nature 371(6494):215–220.

9. Zurovcova M, Eanes WF (1999) Lack of nucleotide polymorphism in the Y-linked sperm flagellar dynein gene Dhc-Yh3 of Drosophila melanogaster and D. simulans. Genetics 153(4):1709–1715.

10. Bachtrog D (2004) Evidence that positive selection drives Y-chromosome degeneration in Drosophila miranda. Nat Genet 36(5):518–522.

11. Chippindale AK, Rice WR (2001) Y chromosome polymorphism is a strong determinant of male fitness in Drosophila melanogaster. Proc Natl Acad Sci USA 98(10):5677–5682.

12. Rohmer C, David JR, Moreteau B, Joly D (2004) Heat induced male sterility in Drosophila melanogaster: adaptive genetic variations among geographic populations and role of the Y chromosome. J Exp Biol 207(Pt 16):2735–2743.

13. Griffin RM, Le Gall D, Schielzeth H, Friberg U (2015) Within-population Y-linked genetic variation for lifespan in Drosophila melanogaster. J Evol Biol 28(11):1940–1947.

14. Lemos B, Araripe LO, Hartl DL (2008) Polymorphic Y chromosomes harbor cryptic variation with manifold functional consequences. Science 319(5859):91–93.

15. Lemos B, Branco AT, Hartl DL (2010) Epigenetic effects of polymorphic Y chromosomes modulate chromatin components, immune response, and sexual conflict. Proc Natl Acad Sci USA 107(36):15826–15831.

16. Francisco FO, Lemos B (2014) How Do Y-Chromosomes Modulate Genome-Wide Epigenetic States: Genome Folding, Chromatin Sinks, and Gene Expression. Journal of Genomics 2:94–103.

17. Brown EJ, Nguyen AH, Bachtrog D (2020) The Drosophila Y chromosome affects heterochromatin integrity genome-wide. Mol Biol Evol.

18. Lancefield DE (1929) A genetic study of crosses of two races or physiological species ofDrosophila obscura. Zeitschrift für Induktive Abstammungs-und Vererbungslehre 52(1):287–317.

19. Dobzhansky T (1937) Further Data on the Variation of the Y Chromosome in Drosophila Pseudoobscura. Genetics 22(3):340–346.

20. Clark AG (1990) Two tests of Y chromosomal variation in male fertility of Drosophila melanogaster. Genetics 125(3):527–534.

21. Bracewell R, Chatla K, Nalley MJ, Bachtrog D (2019) Dynamic turnover of centromeres drives karyotype evolution in Drosophila. Elife 8:923.

22. Johnston JS, Schoener M, Mcmahon D (2013) DNA Underreplication in the Majority of Nuclei in the Drosophila Melanogaster Thorax: Evidence from Suur and Flow Cytometry. Journal of Molecular Biology Research 3(1):47–54.

23. Saksouk N, Simboeck E, Déjardin J (2015) Constitutive heterochromatin formation and transcription in mammals. Epigenetics Chromatin 8:3.

24. Bosco G, Campbell P, Leiva-Neto JT, Markow TA (2007) Analysis of Drosophila species genome size and satellite DNA content reveals significant differences among strains as well as between species. Genetics 177(3):1277–1290.

25. Hoskins RA, et al. (2007) Sequence finishing and mapping of Drosophila melanogaster heterochromatin. Science 316(5831):1625–1628.

26. Chang C-H, Larracuente AM (2019) Heterochromatin-Enriched Assemblies Reveal the Sequence and Organization of the Drosophila melanogaster Y Chromosome. Genetics 211(1):333–348.

27. Koren S, et al. (2017) Canu: scalable and accurate long-read assembly via adaptive k-mer weighting and repeat separation. Genome Res 27(5):722–736.

28. Chin C-S, et al. (2016) Phased diploid genome assembly with single-molecule real-time sequencing. Nat Methods 13(12):1050–1054.

29. Chakraborty M, Baldwin-Brown JG, Long AD, Emerson JJ (2016) Contiguous and accurate de novo assembly of metazoan genomes with modest long read coverage. Nucleic Acids Res 44(19):e147.

30. Pimpinelli S, et al. (1995) Transposable elements are stable structural components of Drosophila melanogaster heterochromatin. Proc Natl Acad Sci USA 92(9):3804–3808.

31. Carvalho AB, Clark AG (2005) Y chromosome of D. pseudoobscura is not homologous to the ancestral Drosophila Y. Science 307(5706):108–110.

32. Bracewell R, Bachtrog D (2020) Complex evolutionary history of the Y chromosome in flies of the Drosophila obscura species group. Genome Biol Evol. doi:10.1093/gbe/evaa051.

33. Goubert C, et al. (2015) De novo assembly and annotation of the Asian tiger mosquito (Aedes albopictus) repeatome with dnaPipeTE from raw genomic reads and comparative analysis with the yellow fever mosquito (Aedes aegypti). Genome Biol Evol 7(4):1192–1205.

34. Hill T, Betancourt AJ (2018) Extensive exchange of transposable elements in the Drosophila pseudoobscura group. Mob DNA 9(1):20.

35. Pritham EJ, Feschotte C (2007) Massive amplification of rolling-circle transposons in the lineage of the bat Myotis lucifugus. Proc Natl Acad Sci USA 104(6):1895–1900.

36. McKee BD, Habera L, Vrana JA (1992) Evidence that intergenic spacer repeats of Drosophila melanogaster rRNA genes function as X-Y pairing sites in male meiosis, and a general model for achiasmatic pairing. Genetics 132(2):529–544.

37. Larracuente AM, Noor MAF, Clark AG (2010) Translocation of Y-linked genes to the dot chromosome in Drosophila pseudoobscura. Mol Biol Evol 27(7):1612–1620.

38. Krupovic M, Koonin EV (2015) Polintons: a hotbed of eukaryotic virus, transposon and plasmid evolution. Nat Rev Microbiol 13(2):105–115.

39. Makałowski W, Gotea V, Pande A, Makałowska I (2019) Transposable Elements: Classification, Identification, and Their Use As a Tool For Comparative Genomics. Methods Mol Biol 1910:177–207.

40. Brown EJ, Bachtrog D (2014) The chromatin landscape of Drosophila: comparisons between species, sexes, and chromosomes. Genome Res 24(7):1125–1137.

41. Brown EJ, Nguyen AH, Bachtrog D (2020) The Y chromosome may contribute to sex-specific ageing in Drosophila. Nat Ecol Evol 37:466–10.

42. Wei KH-C, Gibilisco L, Bachtrog D (2020) Epigenetic conflict on a degenerating Y chromosome increases mutational burden in Drosophila males. Nat Commun 11(1):5537–9.

43. Nguyen AH, Bachtrog D (2021) Toxic Y chromosome: Increased repeat expression and age-associated heterochromatin loss in male Drosophila with a young Y chromosome. PLoS Genet 17(4):e1009438.

44. Sun L, Yu R, Dang W (2018) Chromatin Architectural Changes during Cellular Senescence and Aging. Genes (Basel) 9(4):211.

45. Dobzhansky T, Boche RD (1933) Intersterile races of Drosophila pseudoobscura. Biol Zentr 53:314–330.

46. Dobzhansky T (1933) On the Sterility of the Interracial Hybrids in Drosophila Pseudoobscura. Proc Natl Acad Sci USA 19(4):397–403.

47. Chakraborty BM, Chakraborty R (1984) Variations in the length of human Y chromosome: a statistical study. Acta Anthropogenet 8(3-4):269–275.

48. Altinordu F, Peruzzi L, Yu Y, He X (2016) A tool for the analysis of chromosomes: KaryoType. Taxon 65(3):586–592.

49. Ellis LL, et al. (2014) Intrapopulation genome size variation in D. melanogaster reflects life history variation and plasticity. PLoS Genet 10(7):e1004522.

50. Linford NJ, Bilgir C, Ro J, Pletcher SD (2013) Measurement of lifespan in Drosophila melanogaster. J Vis Exp (71):e50068.

51. Brind’Amour J, et al. (2015) An ultra-low-input native ChIP-seq protocol for genome-wide profiling of rare cell populations. Nat Commun 6:6033.

52. Kapusta A, Suh A, Feschotte C (2017) Dynamics of genome size evolution in birds and mammals. Proc Natl Acad Sci USA 114(8):E1460–E1469.

53. McGurk MP, Barbash DA (2018) Double insertion of transposable elements provides a substrate for the evolution of satellite DNA. Genome Res 28(5):714–725.

54. Yu G (2020) Using ggtree to Visualize Data on Tree-Like Structures. Curr Protoc Bioinformatics 69 (1):e96.

55. Rambaut, A. FigTree. https://tree.bio.ed.ac.uk/software/figtree/ (2007).

56. Kim D, Paggi JM, Park C, Bennett C, Salzberg SL (2019) Graph-based genome alignment and genotyping with HISAT2 and HISAT-genotype. Nat Biotechnol 37(8):907–915.

57. Liao Y, Smyth GK, Shi W (2014) featureCounts: an efficient general purpose program for assigning sequence reads to genomic features. Bioinformatics 30(7):923–930.

58. Love MI, Huber W, Anders S (2014) Moderated estimation of fold change and dispersion for RNA-seq data with DESeq2. Genome Biol 15(12):550–21.

59. Jin Y, Tam OH, Paniagua E, Hammell M (2015) TEtranscripts: a package for including transposable elements in differential expression analysis of RNA-seq datasets. Bioinformatics 31(22):3593–3599.

60. Li H, Durbin R (2009) Fast and accurate short read alignment with Burrows-Wheeler transform. Bioinformatics 25(14):1754–1760.

