## Supplementary Figures for "Transposable element accumulation drives size differences among polymorphic Y chromosomes in Drosophila"

Supplementary figures for
A.H.Nguyen & D. Bachtrog 2021

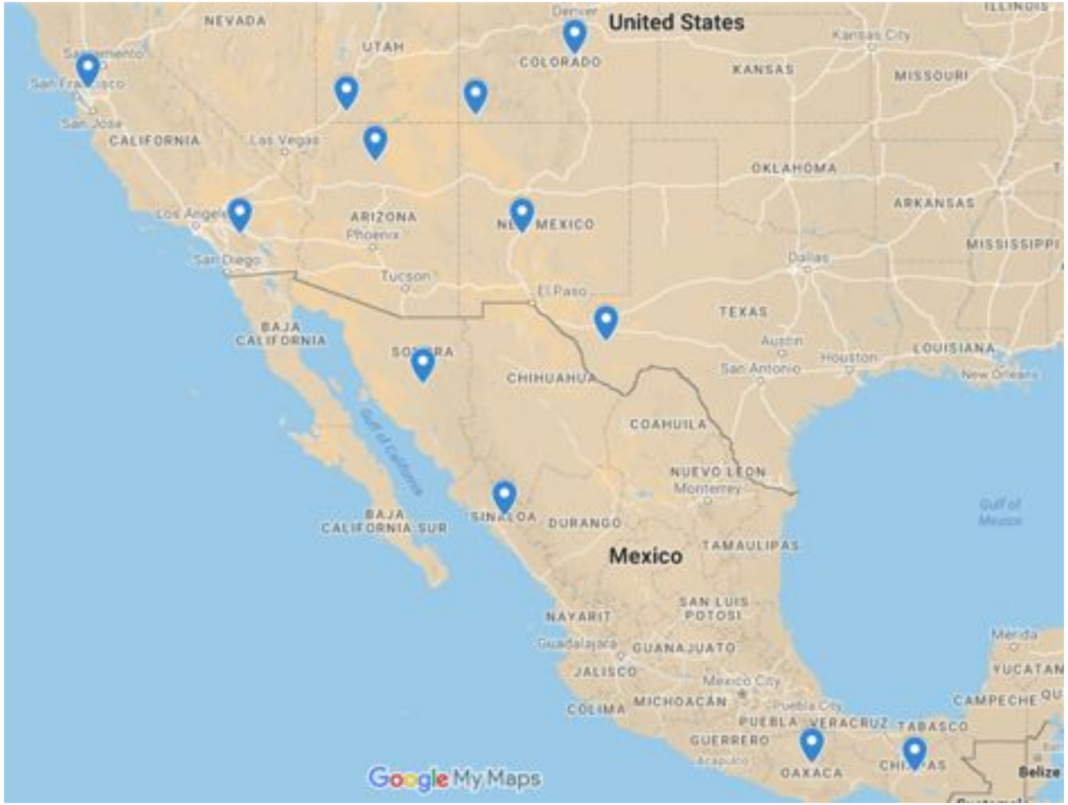


**Figure S1. Geographic locations of sampled D. pseudoobscura Y chromosomes.**

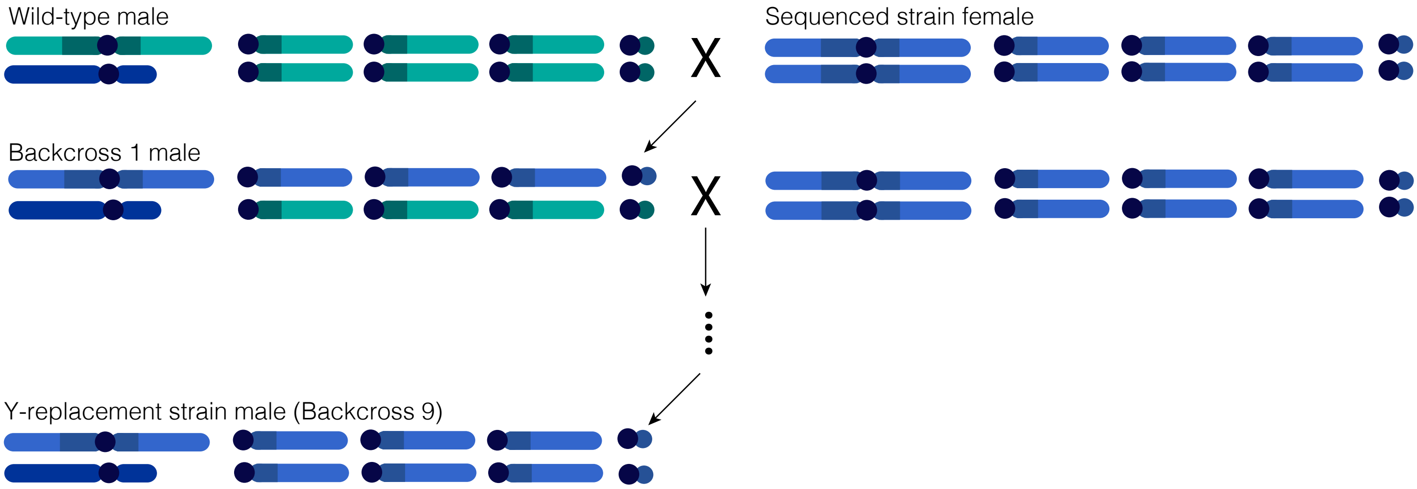


**Figure S2. Backcrossing scheme to generate Y-replacement lines.**

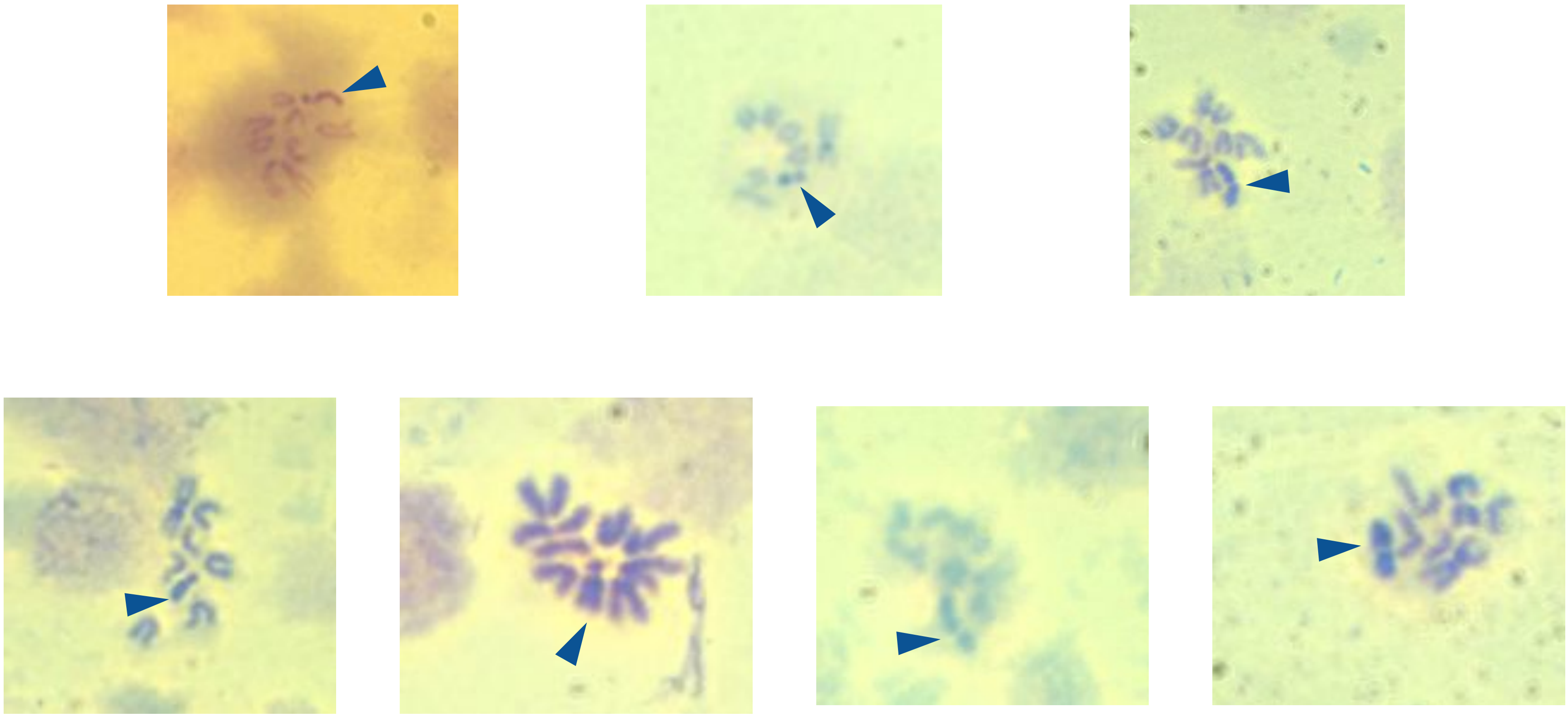

Figure S3. Chromosome squashes. Top (L-R): telocentric, submetacentric, and metacentric Ys. Bottom (L-R): Y-replacement lines with diploid genome sizes of 316Mb, 322Mb, 332Mb, and 346Mb. Arrowhead denotes Y.

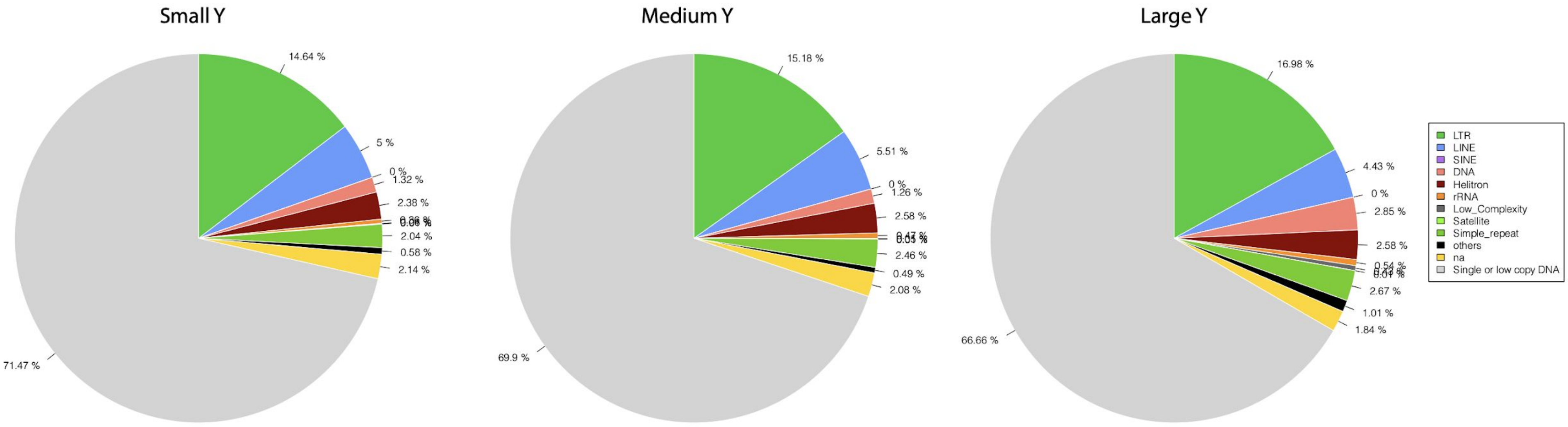


**Figure S4. Reference-free total TE abundance from DNAPipeTE.**

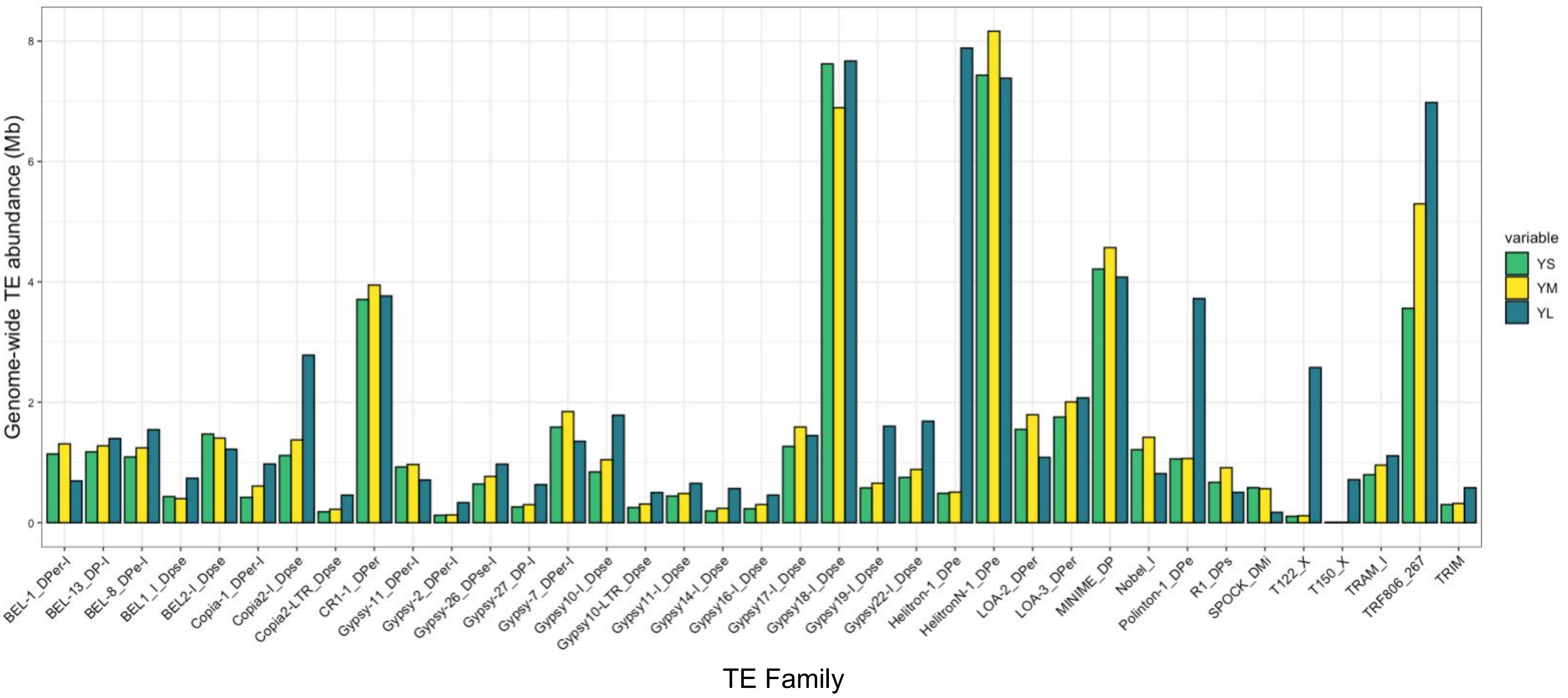


**Figure S5. Total TE abundance from Illumina mappings to libraries for TEs with at least 200kb difference.**

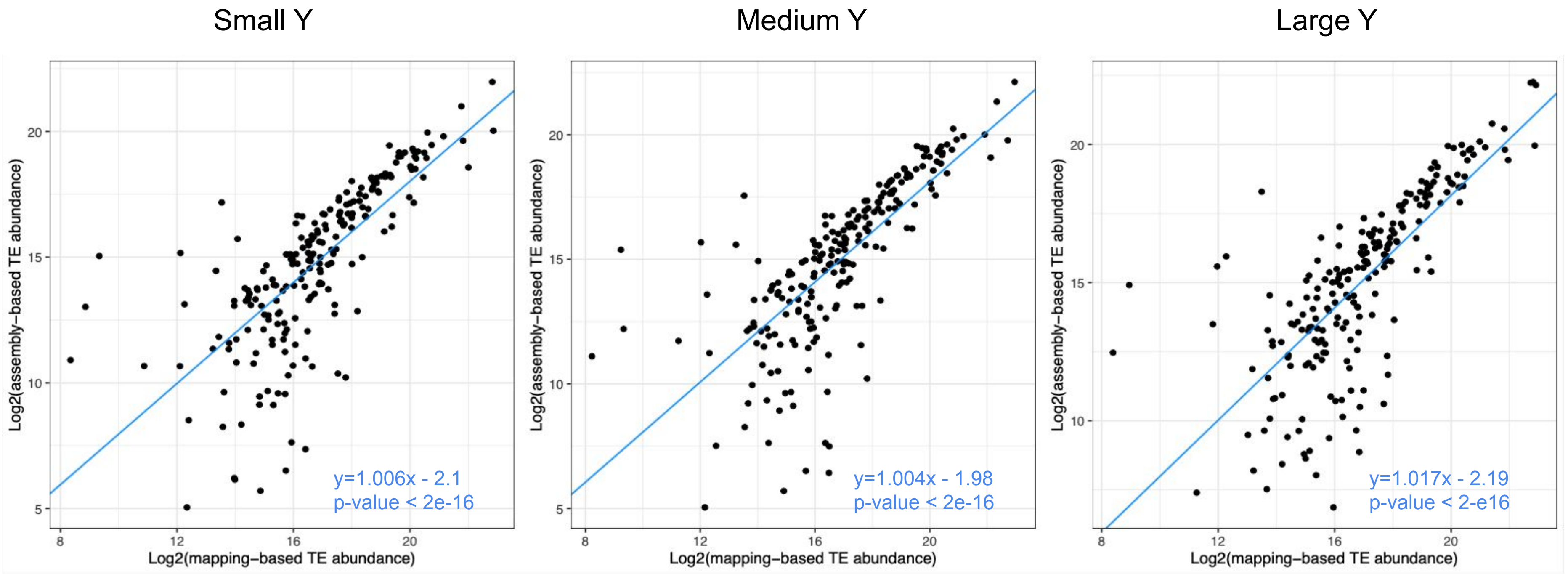


**Figure S6. Correlation between mapping-based and assembly-based TE abundances.**

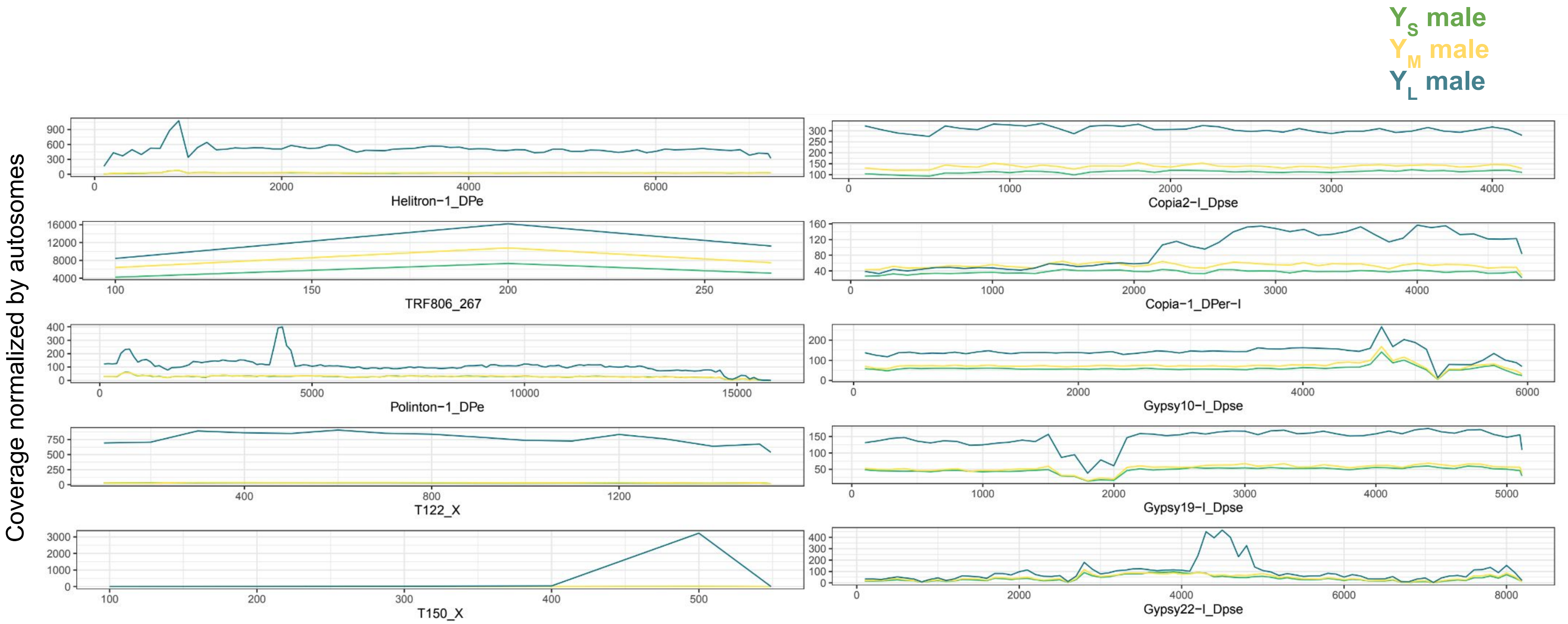


**Figure S7. (A) Coverage of top 10 most abundant transposons in $Y_L$ compared to $Y_S$ from Illumina mappings.**

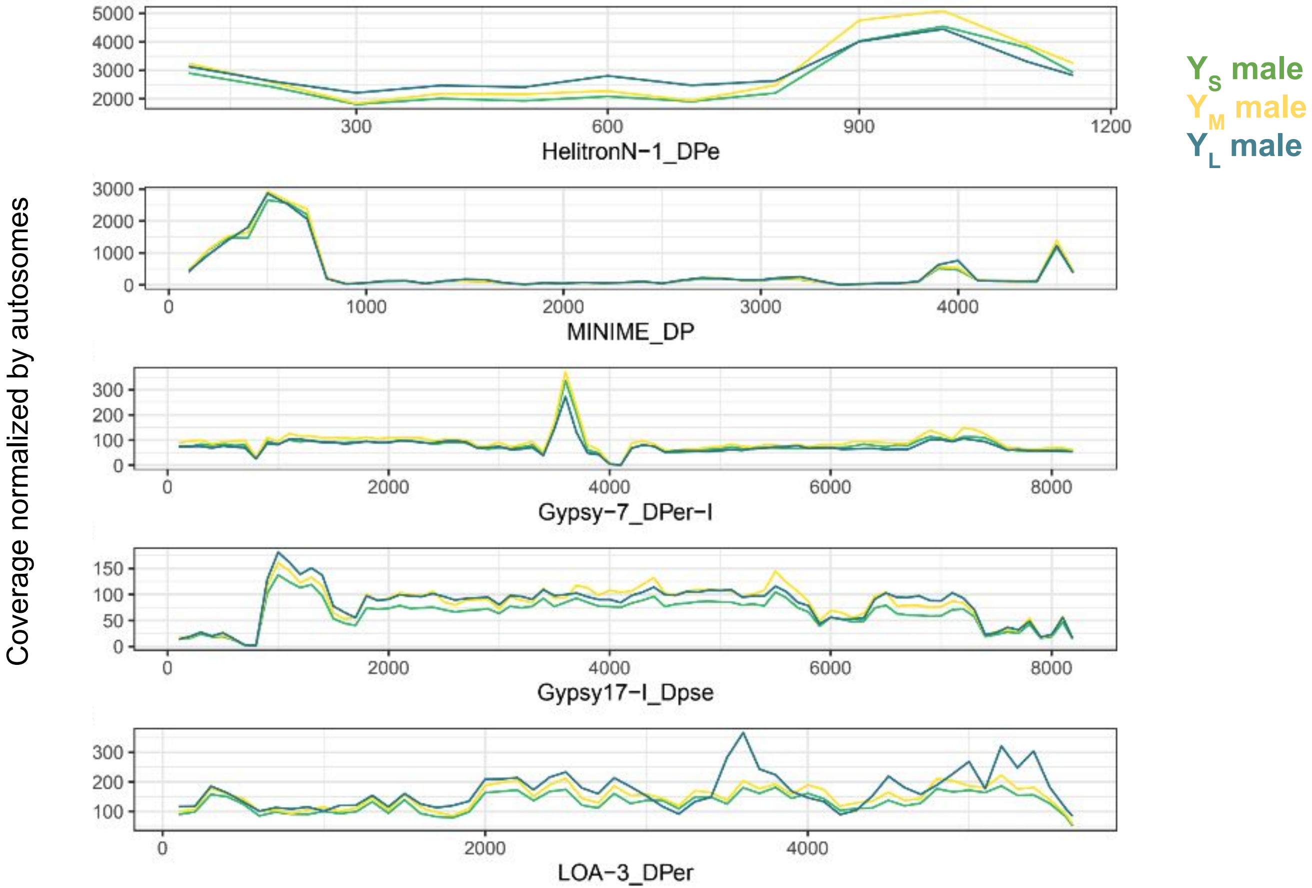


**Figure S7. (B) Coverage of top 5 most abundant transposons in $Y_M$ compared to $Y_S$ from Illumina mappings.**

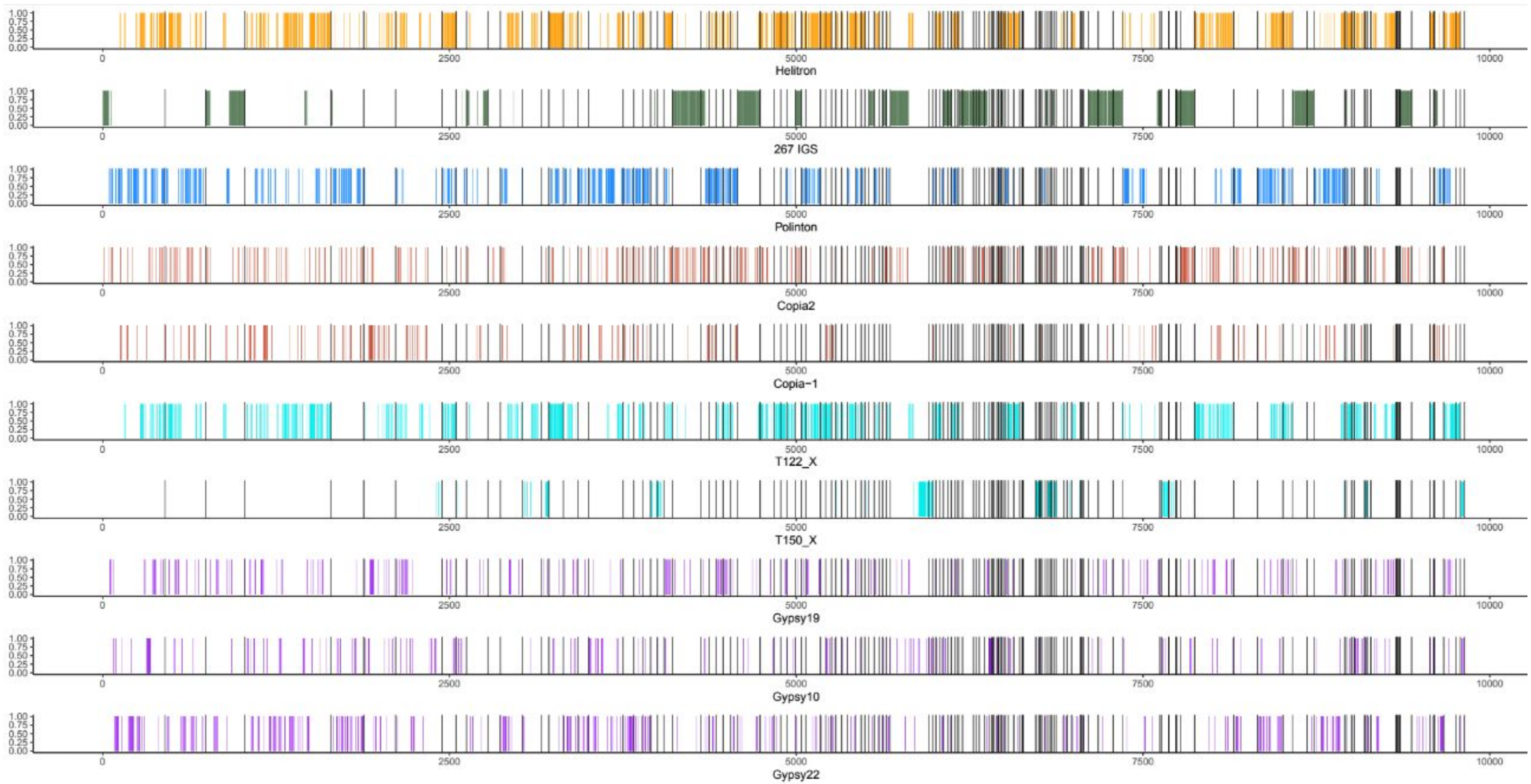


**Figure S8. (A) TE Distribution on Y-linked scaffolds for YL. 1 unit = 1-5kb window.**

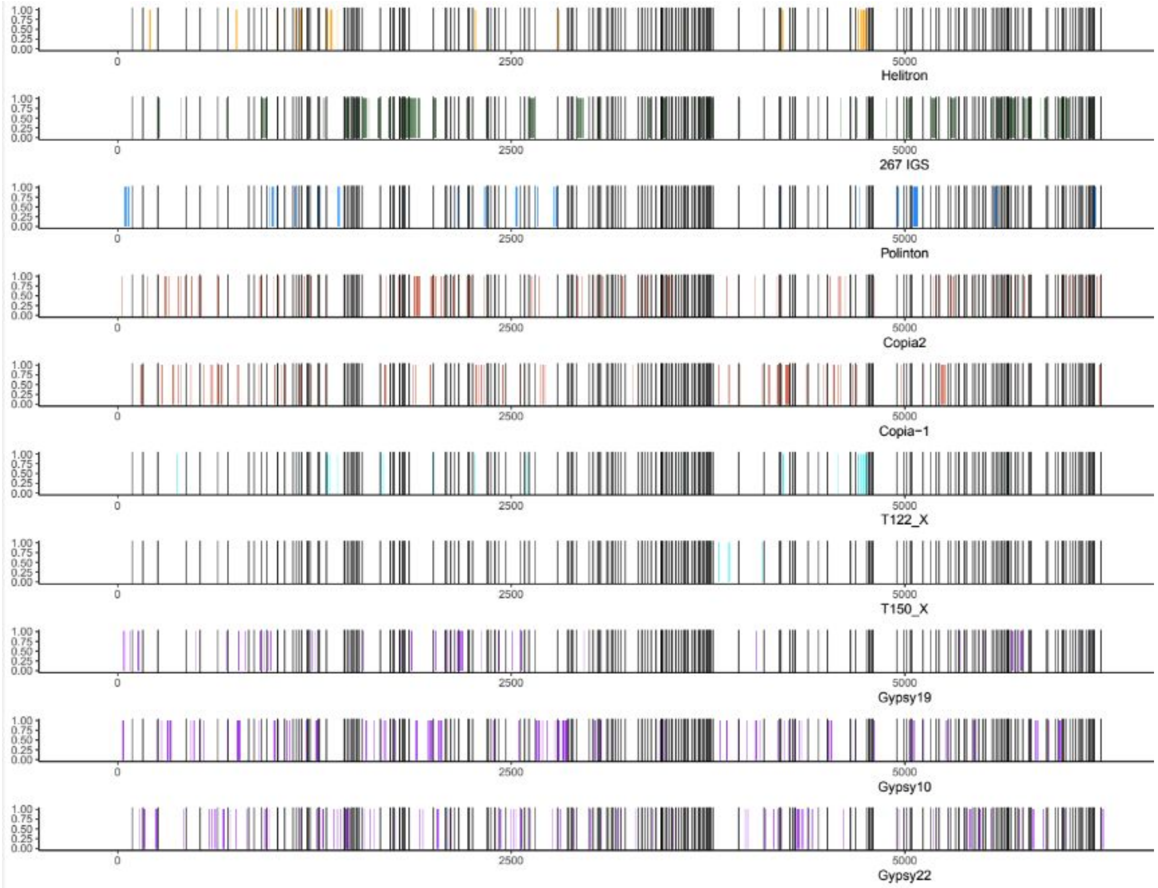


**Figure S8. (B) TE Distribution on Y-linked scaffolds for YM. 1 unit = 1-5kb window.**

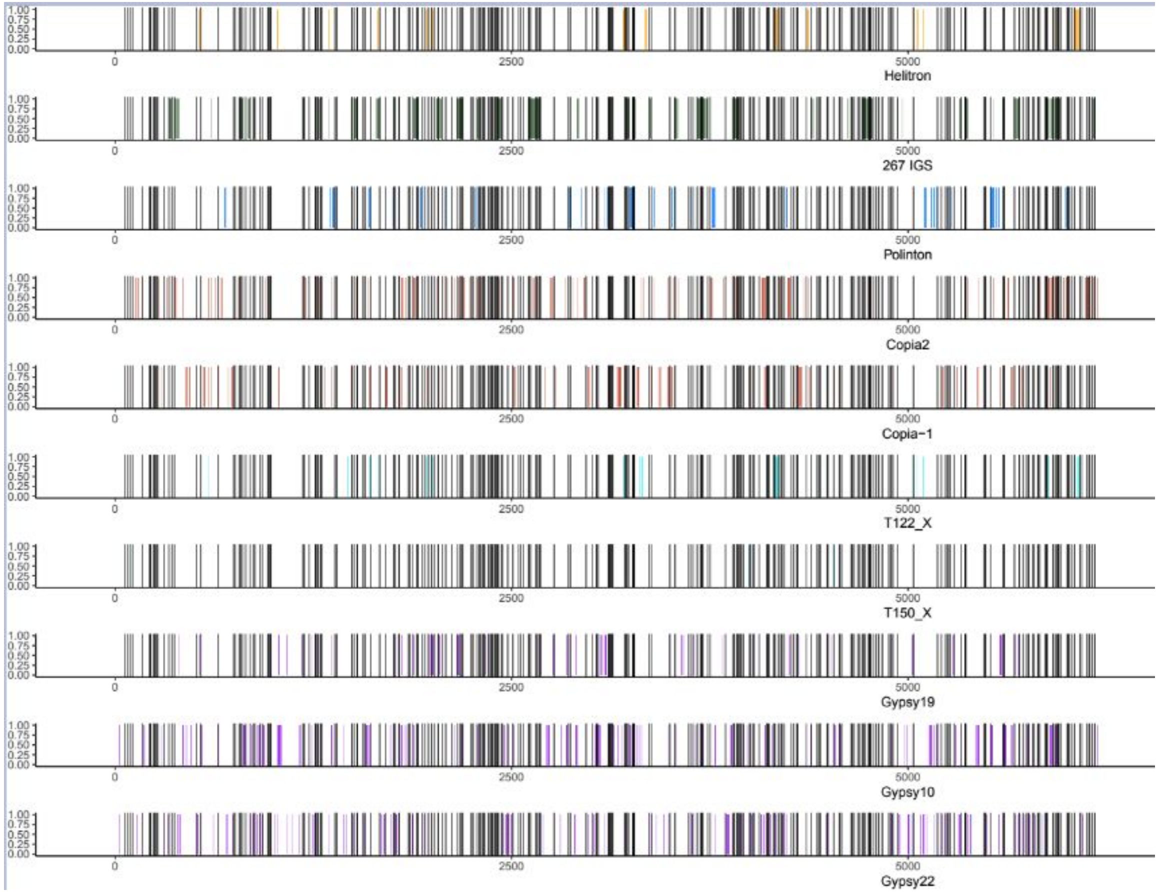


**Figure S8. (C) TE Distribution on Y-linked scaffolds for YS. 1 unit = 1-5kb window.**

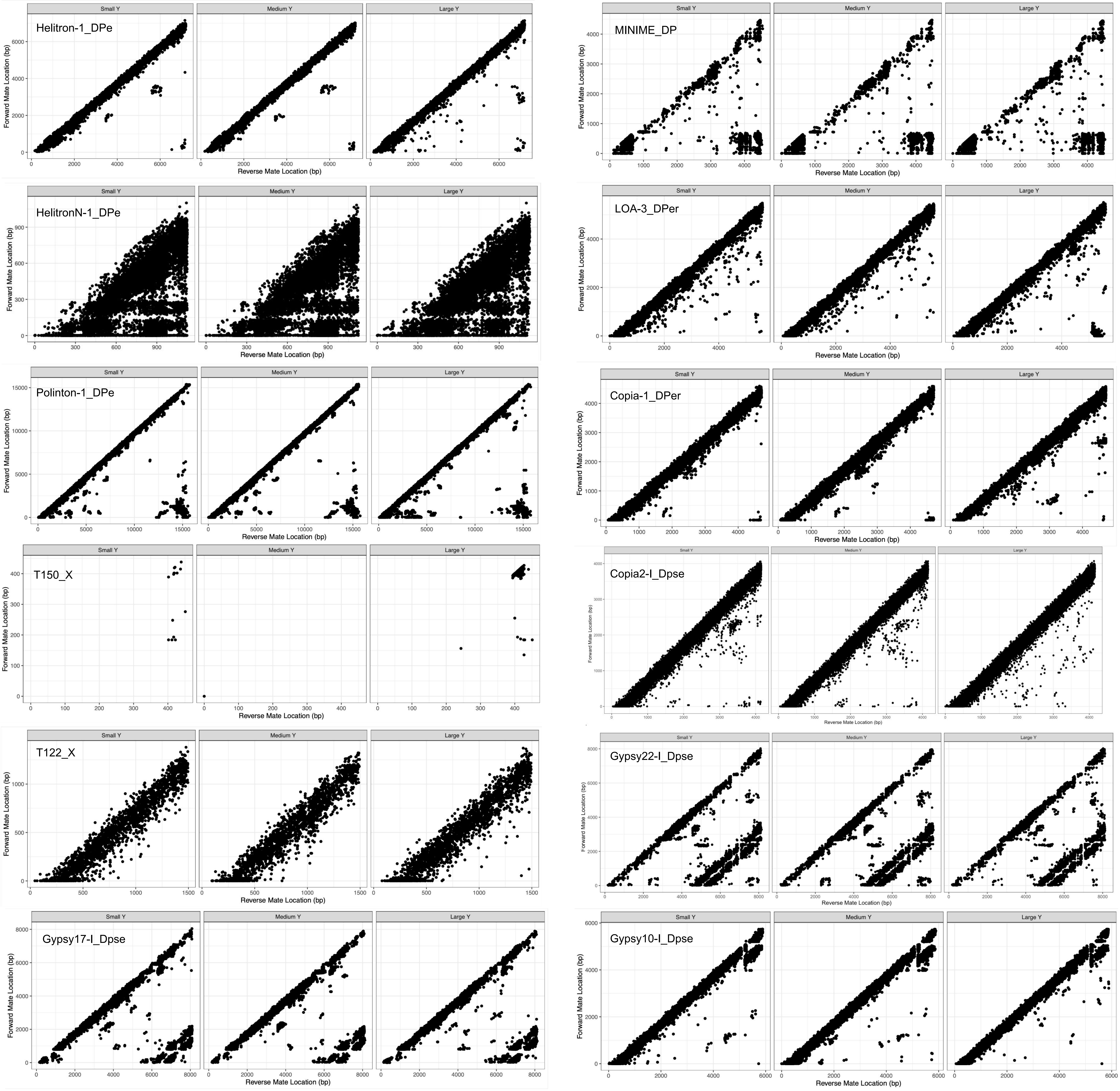


**Figure S9. Inferring tandemly repeated TEs using violations of forward and reverse mapping coordinates.**

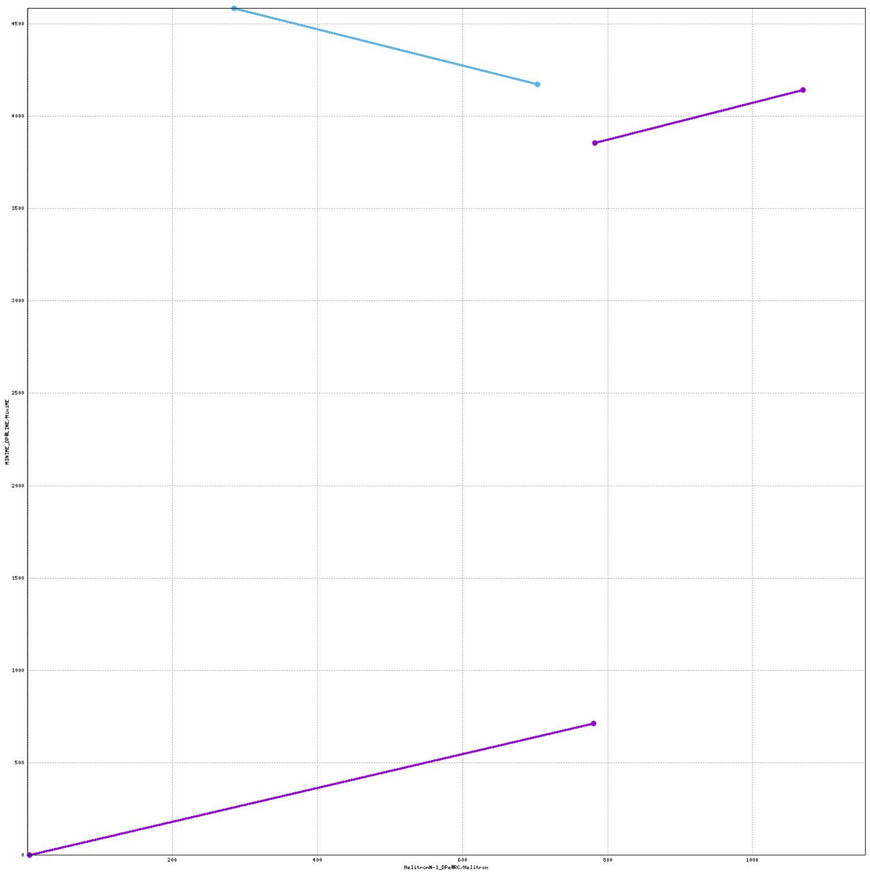


**Figure S10. Alignment of HelitronN-1_DP (X) to MINIME_DP (Y) using nucmer.**

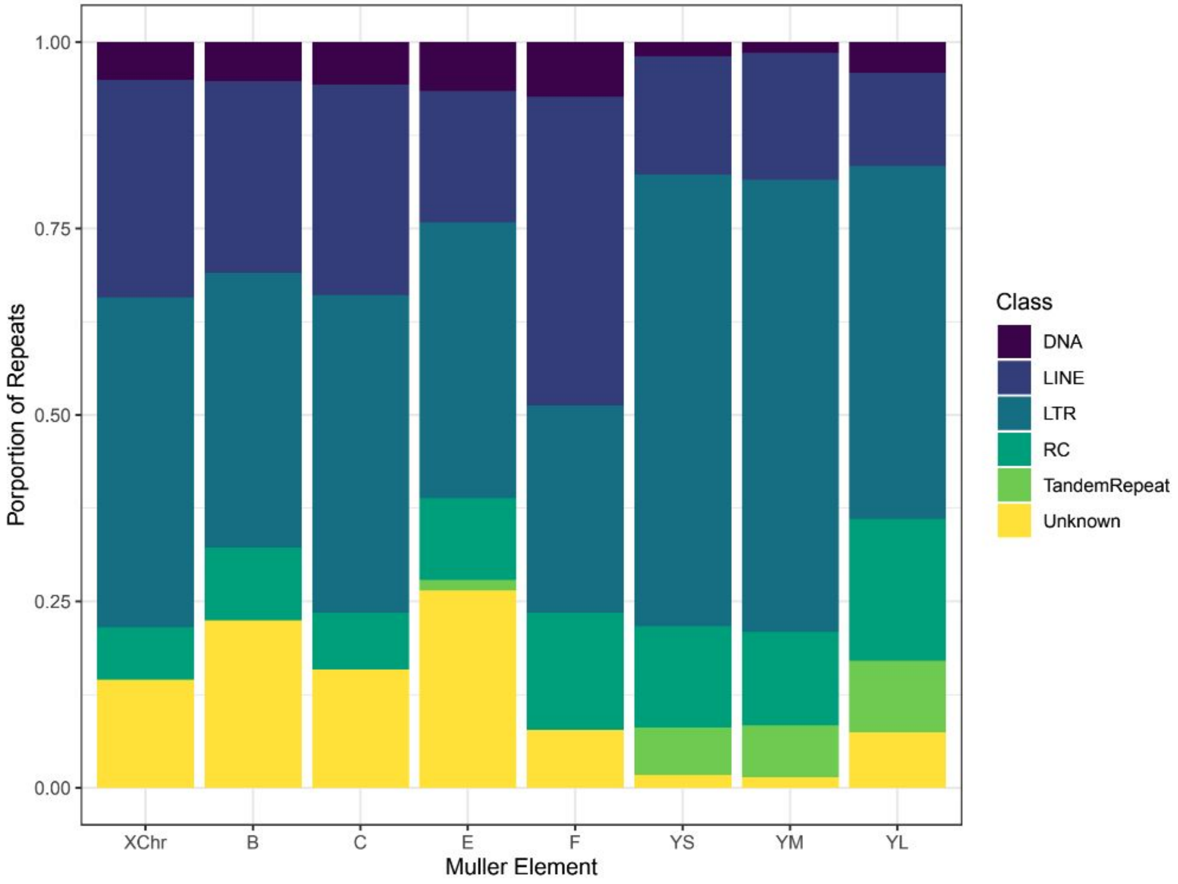


**Figure S11. Repeat composition on a per chromosome basis.**

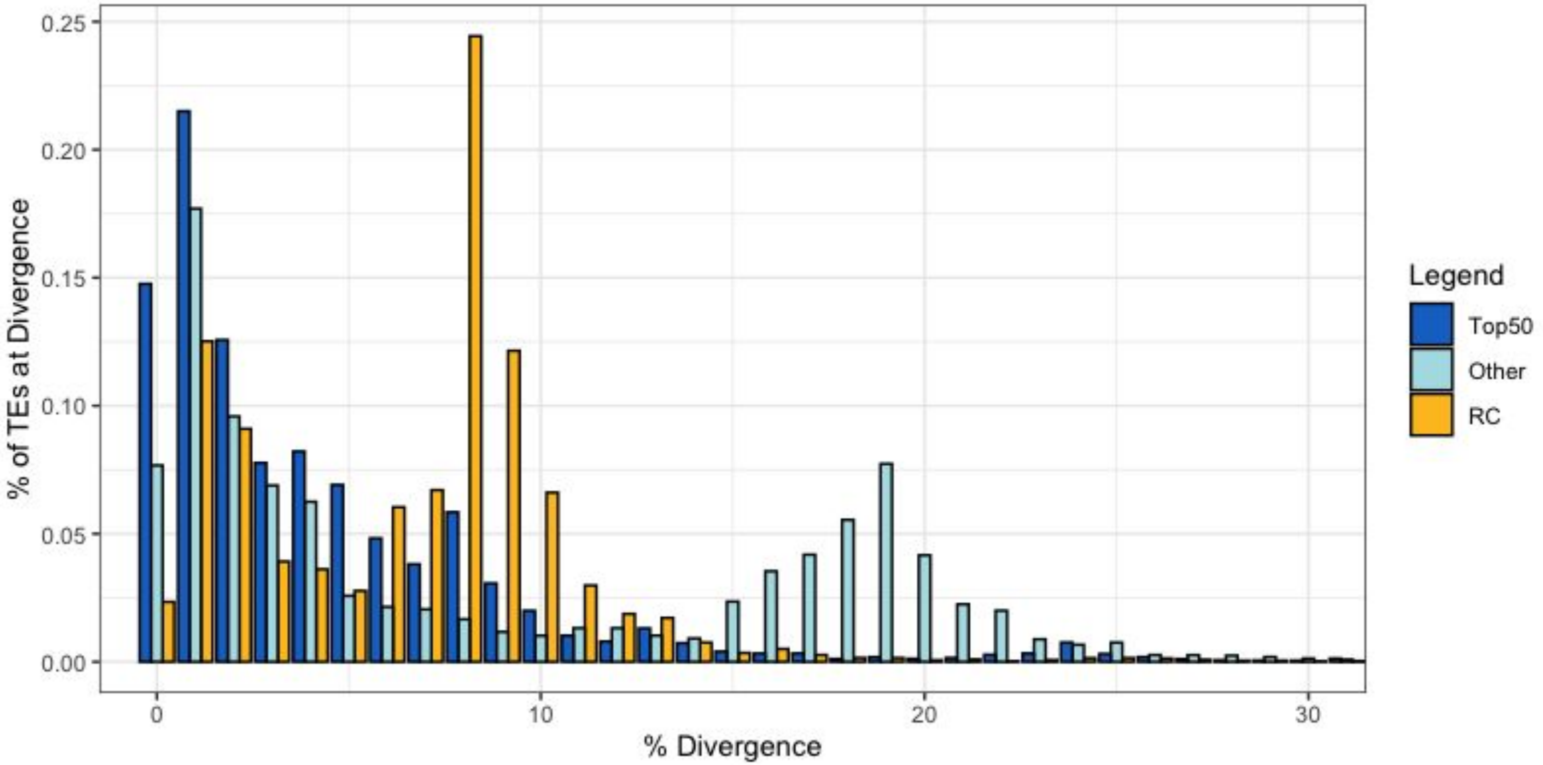


**Figure S12. Divergence of abundant/non-abundant TEs on $Y_M$ relative to $Y_S$.**

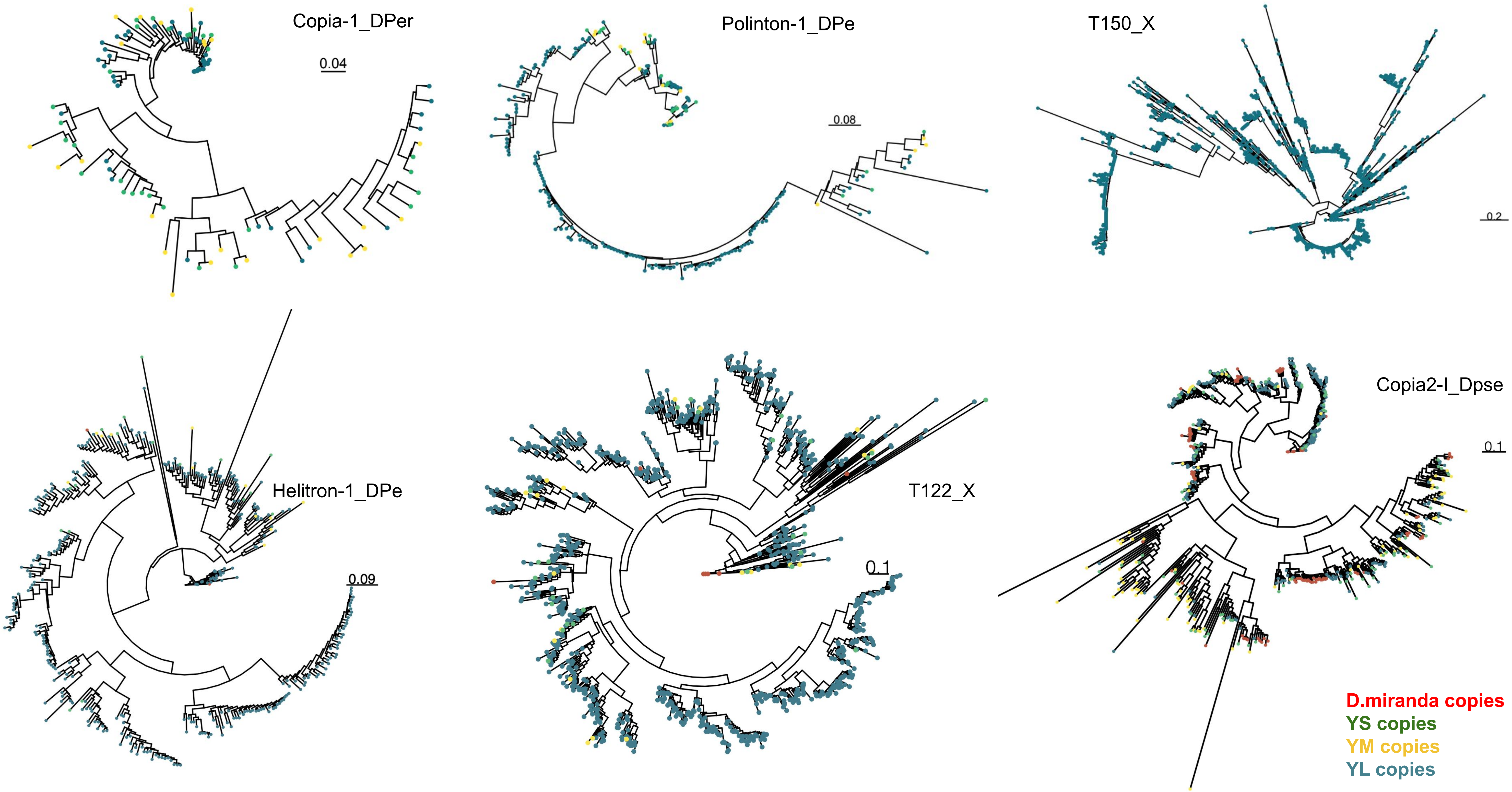


**Figure S13. Unrooted phylogenies of TE copies found on YS, YM, and YL.**

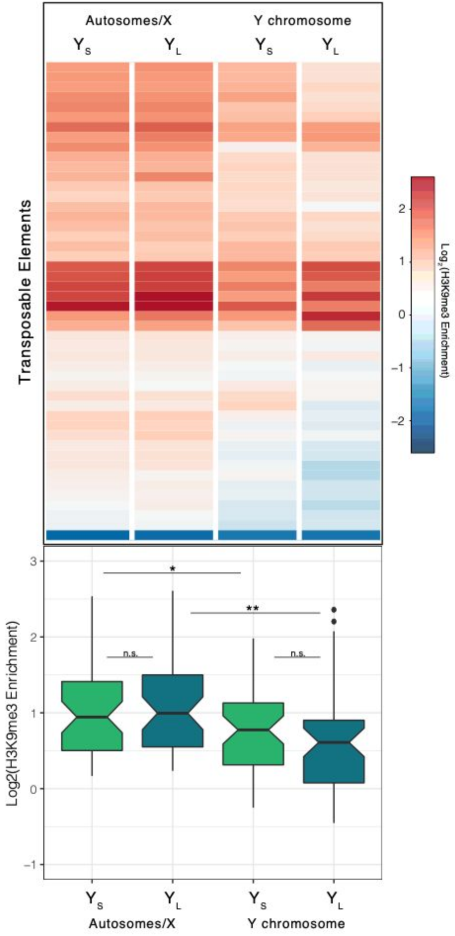


**Figure S14. H3K9me3 enrichment for TEs with 50% higher copy number in either $Y_L$ or $Y_S$.**

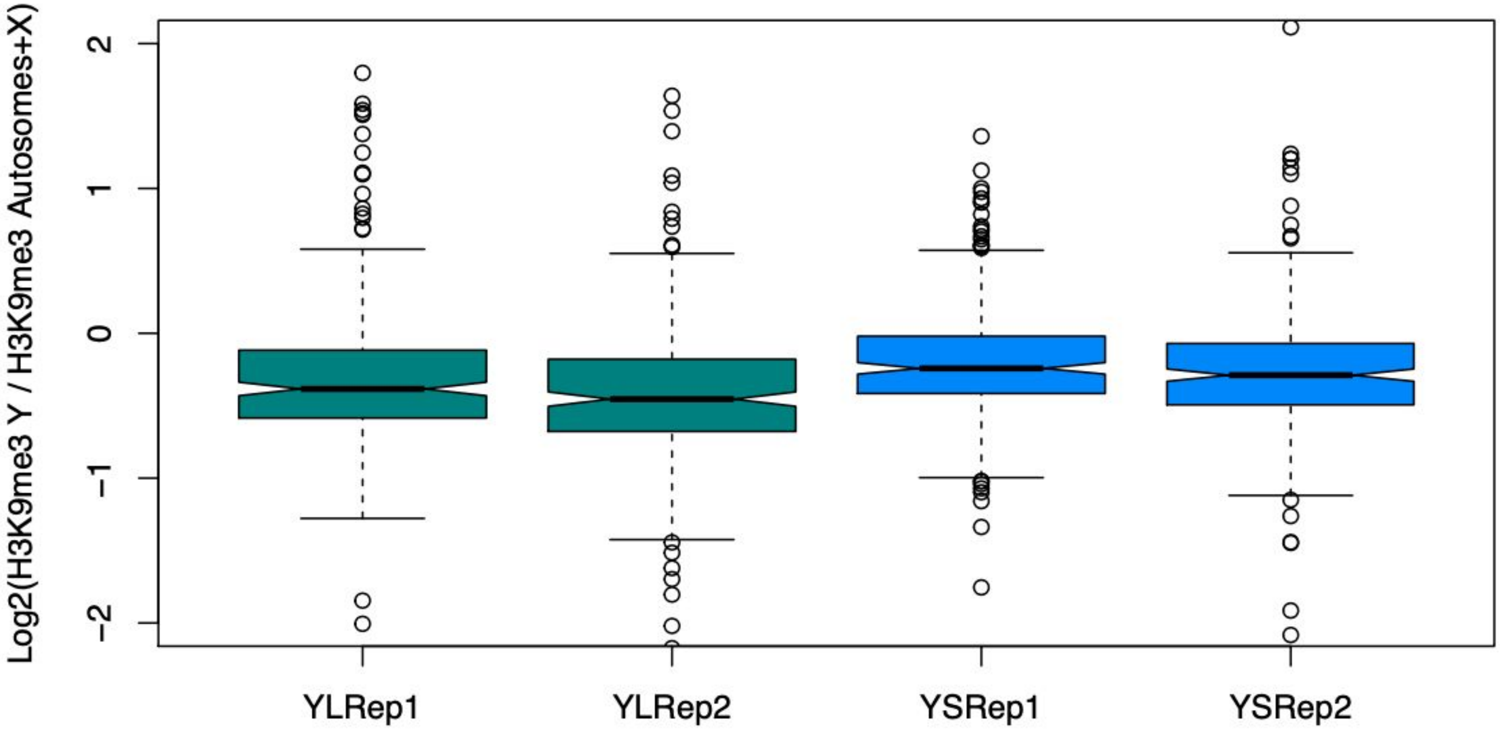


**Figure S15. Log2(H3K9me3 $TE_Y$ / H3K9me3 $TE_{Auto/X}$) for each ChIP replicate.**

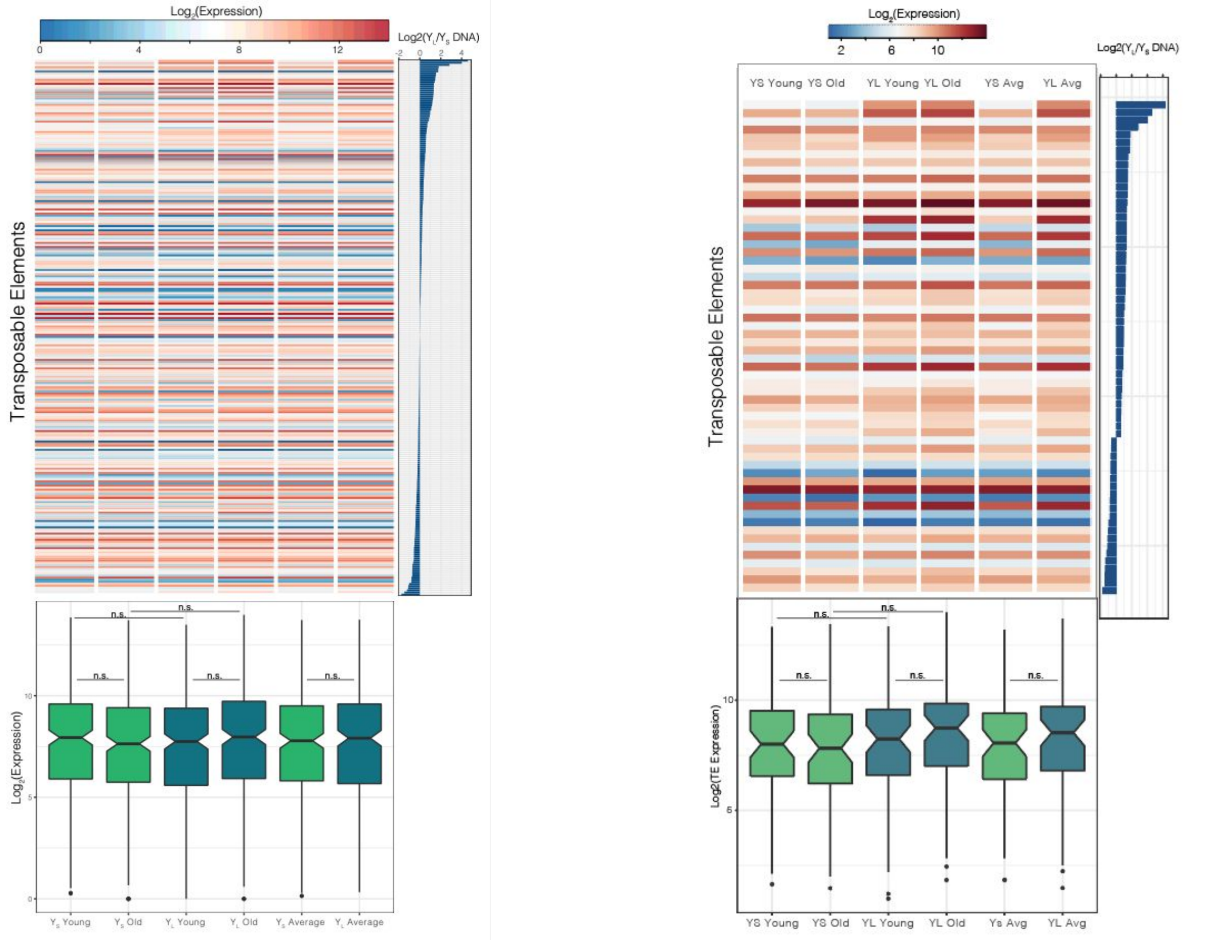


**Figure S16. TE expression by Y-replacement line and age, sorted by TE abundance. (A) All TEs with expression > 1, sorted by Log2(DNA). (B) All TEs with expression > 1 and absolute (Log2(DNA)) > 0.58.**

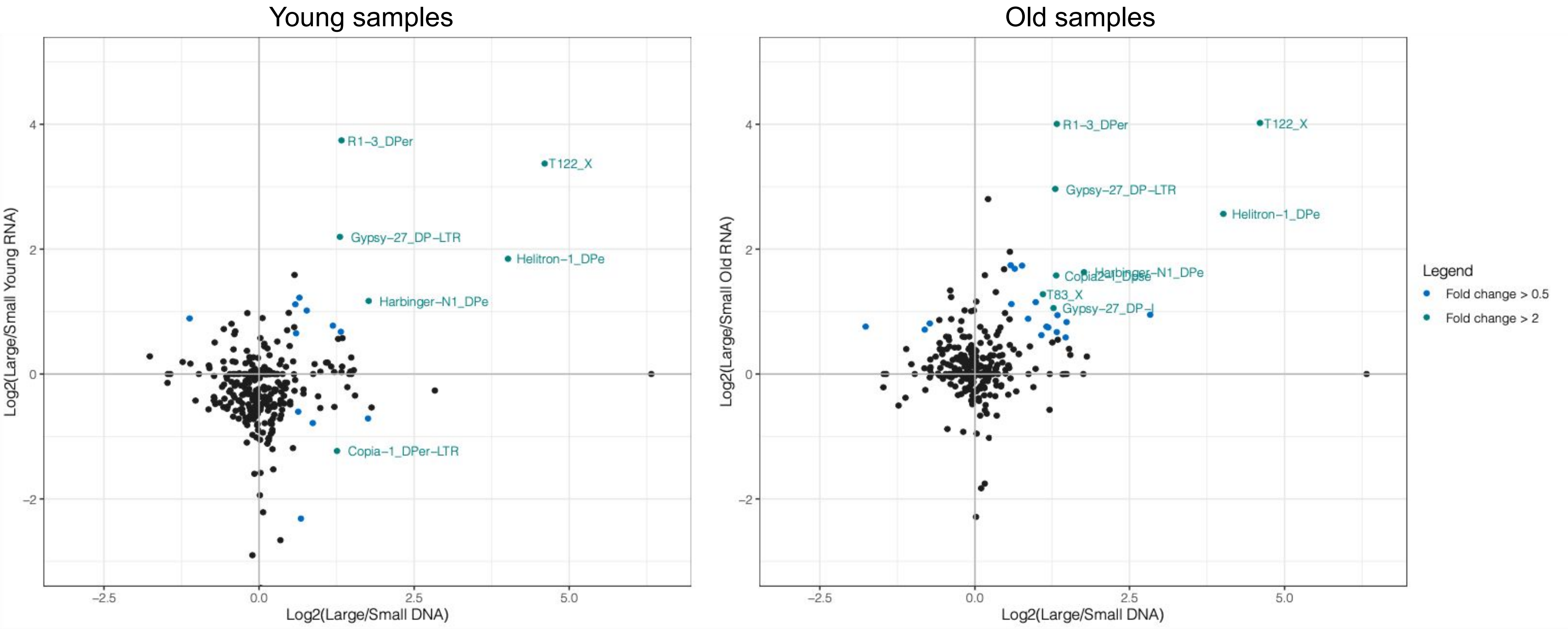


**Figure S17. TE abundance differences vs TE expression differences in Y-replacement line males by age.**

Small Y-replacement line males

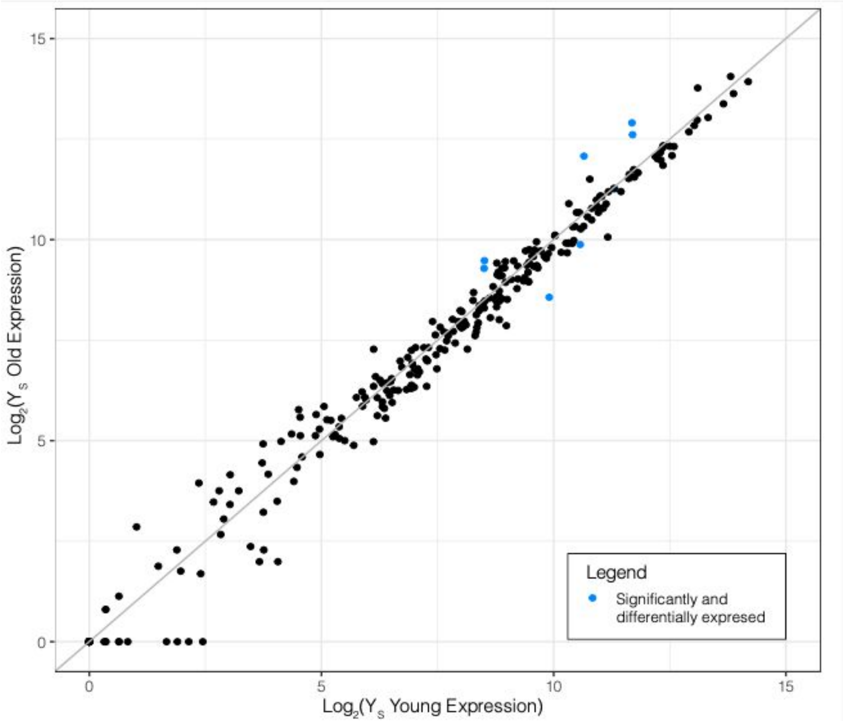


Large Y-replacement line males

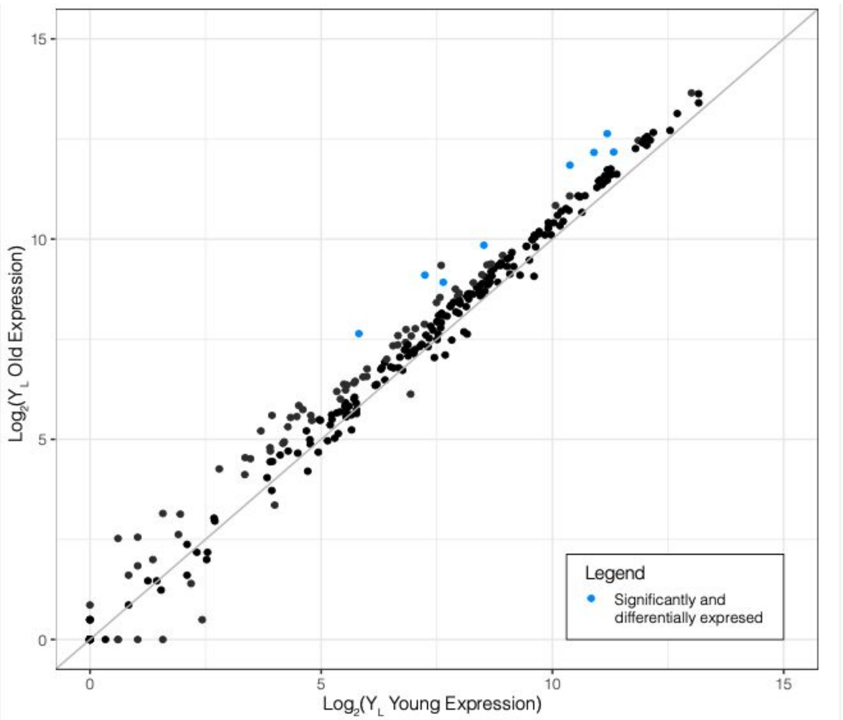


**Figure S18. Age-associated TE expression differences within Y-replacement line males.**

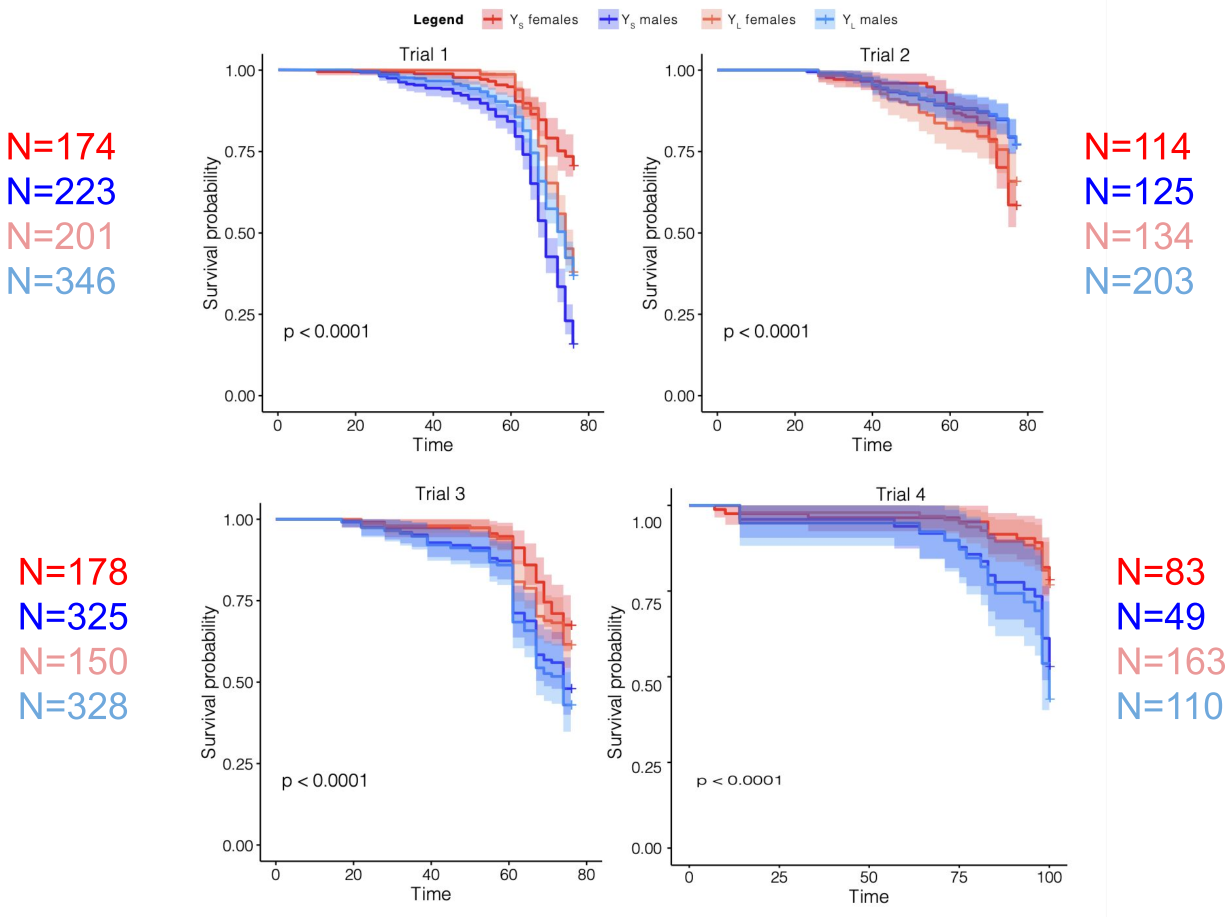


**Figure S19. Y-replacement line aging trials recorded separately.**
